# SH2 domain protein E (SHE) and ABL signaling regulate blood vessel size

**DOI:** 10.1101/2023.07.03.547455

**Authors:** Jennifer A. Schumacher, Zoë A. Wright, Diandra Rufin Florat, Surendra K. Anand, Manish Dasyani, Laurita Klimkaite, Nina O. Bredemeier, Suman Gurung, Gretchen M. Koller, Kalia N. Aguera, Griffin P. Chadwick, Riley D. Johnson, George E. Davis, Saulius Sumanas

## Abstract

Blood vessels in different vascular beds vary in lumen diameter, which is essential for their function and fluid flow along the vascular network. Molecular mechanisms involved in the formation of a vascular lumen of appropriate size, or tubulogenesis, are still only partially understood. *Src homology 2 domain containing E (She)* protein was previously identified in a screen for proteins that interact with Abelson (Abl)-kinase. However, its biological role has remained unknown. Here we demonstrate that She and Abl signaling regulate vascular lumen size in zebrafish embryos and human endothelial cell culture. Zebrafish *she* mutants displayed increased endothelial cell number and enlarged lumen size of the dorsal aorta (DA) and defects in blood flow. Vascular endothelial specific overexpression of *she* resulted in a reduced diameter of the DA lumen, which correlated with the reduced arterial cell number and lower endothelial cell proliferation. Chemical inhibition of Abl signaling in zebrafish embryos caused a similar reduction in the DA diameter and alleviated the *she* mutant phenotype, suggesting that She acts as a negative regulator of Abl signaling. Enlargement of the DA lumen in *she* mutants correlated with an increased endothelial expression of *claudin 5a* and *5b (cldn5a / cldn5b*), which encode proteins enriched in tight junctions. Inhibition of *cldn5a* expression partially rescued the enlarged DA in *she* mutants, suggesting that She regulates DA lumen size, in part, by promoting *cldn5a* expression. SHE knockdown in human endothelial umbilical vein cells resulted in a similar increase in the diameter of vascular tubes, and also increased phosphorylation of a known ABL downstream effector CRKL. These results argue that SHE functions as an evolutionarily conserved inhibitor of ABL signaling and regulates lumen size during vascular tubulogenesis.

## Introduction

Formation of a properly sized vascular lumen, or tubulogenesis, is one of the critical steps during vascular development. Blood vessels vary in size in different vascular beds, which is essential to allow the flow of fluids and blood cells along the vascular network. Various genetic mechanisms have been implicated in establishing and maintaining vascular lumen of a proper size. A family of Rho GTPases which include Rac1, Cdc42 and RhoA have well established roles in regulating cell cytoskeleton rearrangements that take place during vascular tubulogenesis [1–8]. Activity of these Rho GTPases is known to be regulated by integrin-fibronectin signaling [9]. Several other pathways, including VEGF and Notch signaling are also thought to contribute to regulation of vascular lumen size directly [10]. However, our understanding of molecular mechanisms that regulate vascular tubulogenesis is still incomplete.

Abelson (Abl) kinase signaling has diverse roles during morphogenesis such as regulating cytoskeletal organization that is important for cellular protrusions, cell migration, morphogenesis, adhesion, endocytosis and phagocytosis [11]. Chromosomal translocation of human ABL1 to the breakpoint cluster region (BCR) gene results in production of the BCR-ABL1 fusion protein in patients with different types of leukemias [12]. While numerous studies have focused on the role of BCR-ABL1 in leukemias, the role of ABL signaling during developmental morphogenesis is less understood. Intriguingly, Abl signaling is known to regulate cell-cell adhesion through Rho and Rac GTPases. Chemical inhibition of Abl signaling, or mutation of the two ABL homologs Abl1 and Arg in epithelial cell culture, results in activation of RhoA and increased actomyosin contractility, disrupting adherens junctions [13–16] In addition, Abl signaling is known to activate Rac1 in cultured human embryonic kidney cells [17]. Abl signaling is also known to promote cell proliferation in diverse cell types, such as smooth muscle cells and fibroblasts [18, 19]. Previously established roles of Abl signaling in vascular endothelium include regulating angiogenesis, endothelial cell (EC) survival and vascular permeability [20–22]. Recent data also indicate that ABL signaling is increased in a cell culture model of venous malformations (VMs) and that inhibition of ABL signaling reduces lumen diameter in a VM model in vivo [23]. However, the role of ABL signaling in the formation of the vascular lumen during embryogenesis has not been previously investigated.

ABL signaling is known to be regulated by various effector proteins. In a yeast two hybrid screen, two novel Abl-interacting proteins Shd and She were identified [24]. Together with a related Shb protein they comprise a separate family of SH2 proteins. Shd is known to be phosphorylated by Abl kinase in vitro, while She phosphorylation has not been previously investigated [24]. However, biological roles of these proteins remain largely unknown.

We have previously identified *she* as a novel vascular endothelial specific gene, which functions downstream of the ETS transcription factor Etv2 / Etsrp in zebrafish embryos [25]. *She* expression is enriched in arterial vessels, such as the dorsal aorta. Its protein sequence is highly conserved between different vertebrates. However, its biological function has not been previously investigated in any model system.

Here we used genome editing approaches to generate zebrafish *she* genetic mutants. We show that *she* mutants display enlarged dorsal aorta (DA), followed by a failure of circulation and embryonic lethality. In contrast, She overexpression caused a reduction in DA lumen size. Chemical inhibition of Abl signaling caused a similar reduction of the DA size and reversed enlarged aorta observed in *she* mutants. Our data further suggest that She functions in part by regulating expression of *claudin 5a* in the DA. Knockdown of She in human umbilical vein endothelial cells (HUVECs) resulted in a similar dilation of vascular tubes arguing that She function is evolutionarily conserved. In summary, our results identify novel roles for Abl signaling and She in regulating vascular tubulogenesis during embryonic development.

## Results

To determine the function of zebrafish She in cardiovascular development, we used TALEN and CRISPR/Cas9 genome editing to generate two loss-of-function *she* mutant alleles (Suppl. Fig. S1). The *she^ci26^* allele has a 7 bp deletion, which is predicted to result in a frameshift after amino acid 4 and is expected to lead to premature translation termination. The *she^ci30^* allele has a deletion of 575 bp and an insertion of 24 bp which is predicted to lead to a frameshift and a premature stop codon after amino acid 243 (Fig. 1A, Suppl. Fig. S1). *she* mRNA expression in *she^ci26^* mutants was not significantly affected, while *she* mRNA expression in *she^ci30^* mutant embryos was significantly decreased (Fig. 1B-E), indicating a possibility of nonsense-mediated RNA decay.

**Figure 1.**
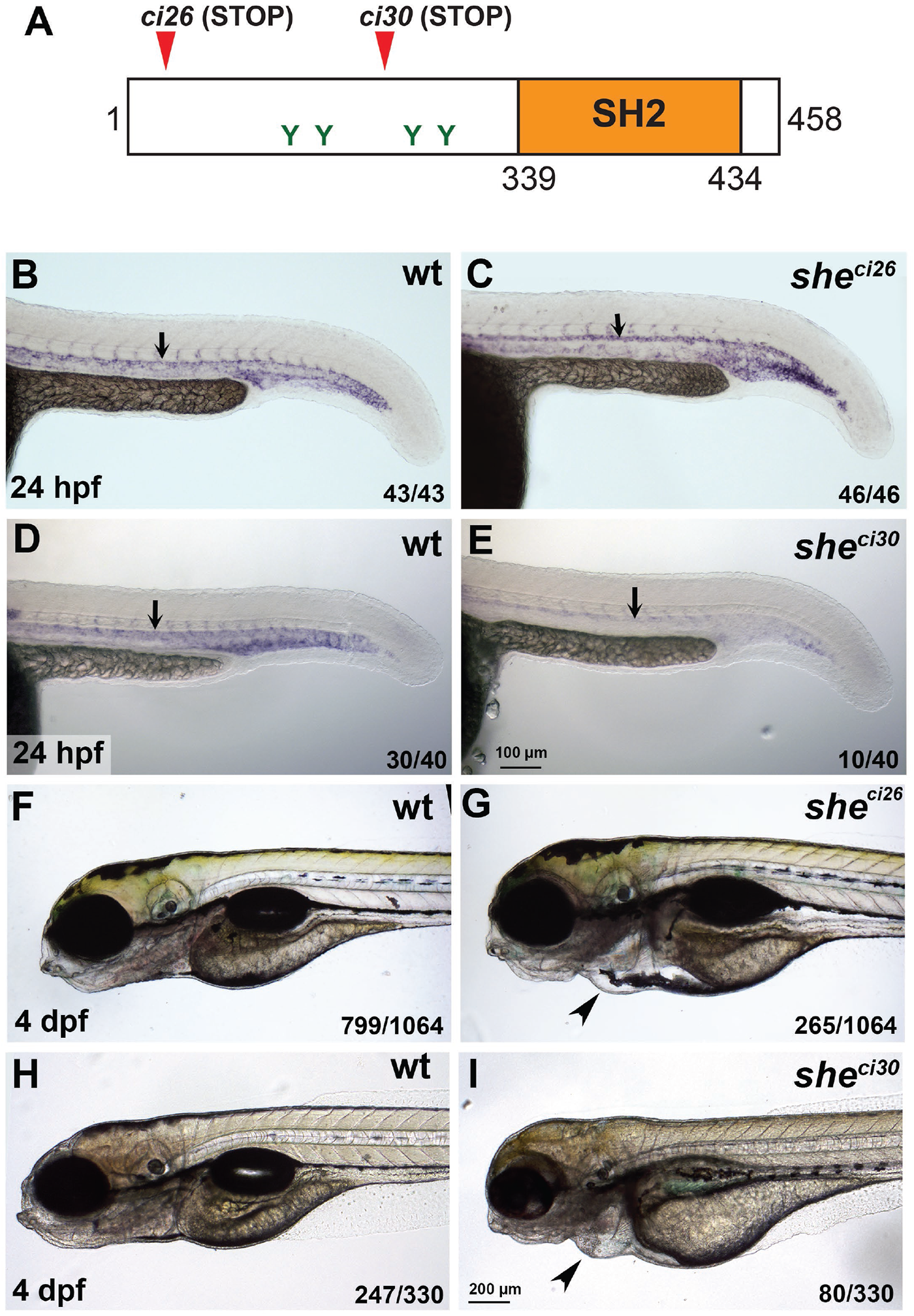
*she* mutants display pericardial edema and loss of blood circulation. (A) Zebrafish SHE protein diagram. *she ci26* and *ci30* mutant alleles are predicted to result in a frameshift and premature stop codons. SH2 domain and consensus tyrosine ABL phosphorylation sites are shown. (B-E) In situ hybridization analysis of *she* expression in *she* mutants at 24 hpf. Note that *she* expression is unaffected in *she^ci26^* embryos while it is strongly reduced in *she^ci30^* embryos. Homozygous *she^ci26^* mutant embryos were obtained by an incross of *she^ci26-/-;^ fli1a:she-2A-mCherry; kdrl:GFP* parents (which are viable due to *fli1a:she-2A-mCherry* rescue) and selected for mCherry negative embryos. wt control embryos were obtained by incross of sibling *wt; fli1a:she-2A-mCherry* parents and selected for mCherry-embryos. *she^ci30^* embryos were obtained by incross of *she^ci30+/-^* parents and genotyped after in situ hybridization. 25% (10 out of 40) embryos showed strong reduction in she expression which correlated with the mutant phenotype. (F-I) Brightfield images of *she^ci26^* and *she^ci30^* mutant embryos and their siblings (wild-type and or heterozygous) at 4 dpf. Note the pericardial edema (arrowheads) in the mutant embryos. Embryos were obtained by the incross of heterozygous parents in *kdrl:GFP* background. 24.9% (265 out of 1064) and 24.2% (80 out of 330) embryos obtained in *she^ci26^* or *she^ci30^* incross, respectively, showed this phenotype. A subset of embryos was genotyped to confirm the correlation between the phenotype and genotype.

Homozygous *she* mutant embryos were then obtained by an incross of respective heterozygous adult carriers and analyzed for morphological defects. The embryos appeared morphologically normal until 4 days-post-fertilization (dpf). At 4-6 dpf both *she^ci26^* and *she^ci30^* mutants developed pericardial edema, and their blood circulation slowed down or completely stopped (Fig. 1F-I, Movies 1,2). As a result, mutant embryos for both alleles were lethal by approximately 7 dpf. *she^ci26^* mutants showed edema and blood circulation failure at a slightly earlier stage (4 dpf) compared to the *she^ci30^* allele (5-6 dpf). *she^ci26^* and *she^ci30^* mutants failed to complement each other, and in the complementation cross 58 out of 262 embryos (22.1%) developed pericardial edema and greatly reduced or absent blood flow by 6 dpf. Because *she^ci26^* allele showed a slightly more severe phenotype, it was used for all further studies.

To analyze potential vascular defects, *she* mutants were crossed into vascular endothelial reporter *kdrl:GFP* background. Overall vascular patterning was normal and unaffected at 1-4 dpf, when analyzed by confocal imaging for GFP expression (Fig. 2A,B). In addition, arterial and venous trunk intersegmental vessel sprouting appeared normal in mutant and wild-type sibling embryos (Suppl. Fig. S2). There was also no effect on endothelial marker *kdrl,* arterial marker *flt1* or venous marker *dab2* expression when analyzed by in situ hybridization at 24 hpf (Suppl. Fig. S3). However, the dorsal aorta (DA) was wider in *she* mutants compared with their sibling wild-type (wt) embryos at both 1 and 2 dpf (Fig. 2C-H,K). Notably, no circulation defects were apparent in *she* mutants at these stages. In contrast, the DA was narrower in *she* mutants at 4 dpf when compared to their siblings (Fig. 2I-K). It is likely that the DA collapse apparent at this stage is a secondary consequence of failing blood circulation. The diameter of the posterior cardinal vein (PCV) was not significantly different between *she* mutant and wild-type embryos at 2-4 dpf, however at 1 dpf, a slight enlargement of the PCV was observed in mutant embryos (Fig. 2K).

**Figure 2.**
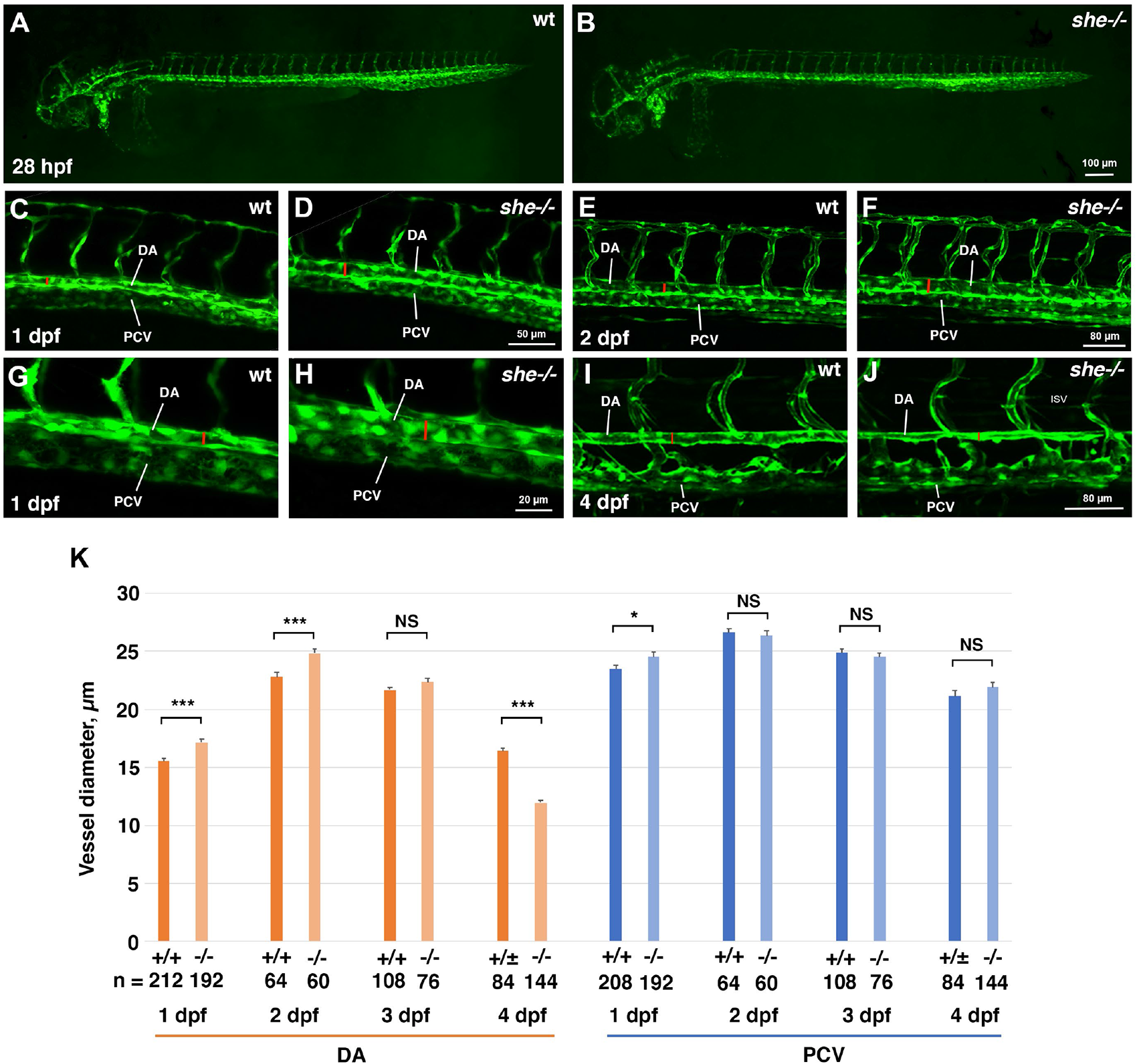
*she* mutants show enlarged diameter of the dorsal aorta. (A,B) Overall vascular patterning of *she-/-* mutants is normal when compared to their sibling wild-type embryos at 28 hpf. Embryos are in *kdrl:GFP* background. (C-H) A wider DA is observed in *she* mutant embryos compared to their wild-type *(she+/+)* siblings at 1 and 2 dpf (28 and 48 hpf respectively). Red line indicates DA diameter. (G,H) shows higher magnification image of trunk vasculature at 1 dpf. (I,J) DA is narrower in *she* mutants at 4 dpf compared to their siblings. (K) Diameter of the DA and PCV at 1-4 dpf in *she* mutants and their wild-type siblings. * p<0.05; ***p<0.001, NS – not significant, Student t-test. Error bars show SEM. At all stages, s*he* mutant and sibling embryos were obtained by in-crossing *she^ci26+/-^*; *kdrl:GFP* carriers. Embryos at 1 and 2 dpf were genotyped after imaging. Embryos at 4 dpf were separated based on the phenotype, and wild-type siblings include *she+/+* and *she+/-* embryos at this stage.

This suggests that She may function to restrict the DA size during vascular development. To test if She function is sufficient to limit the size of the DA, we made a construct which contains the coding sequence for the zebrafish She driven by the vascular endothelial *fli1a* promoter, followed by the self-cleaving peptide 2A and mCherry reporter. Injection of this DNA construct in combination with Tol2 mRNA into zebrafish embryos resulted in mosaic overexpression of *she-2A-mCherry* in vascular endothelial cells. Interestingly, segments of the DA with multiple mCherry-positive cells displayed narrower lumen compared with adjacent mCherry-negative regions of the DA (Fig. 3A,B). A stable zebrafish transgenic reporter line was then established using Tol2-mediated transgenesis and used for further studies (Suppl. Fig. S4A-C). *fli1:she-2A-mCherry* embryos were viable, yet they exhibited a reduction in the DA diameter at 28 hpf, when compared to sibling embryos without the transgene (Fig. 3C,D,I). To test if vascular endothelial *she* expression could rescue the *she* mutant phenotype, the *fli1:she-2A-mCherry* line was crossed into the *she^ci26^* mutant background. Remarkably, *she-/-; fli1:she-2A-mCherry* embryos were viable and had smaller DA diameter when compared with *she-/-* sibling embryos without the transgene (Fig. 3E,F,I). They did not exhibit pericardial edema and showed normal blood circulation at 4 dpf (Fig. 3G,H,J). Altogether, these results show that She functions cell-autonomously in the vascular endothelium to regulate the DA size.

**Figure 3.**
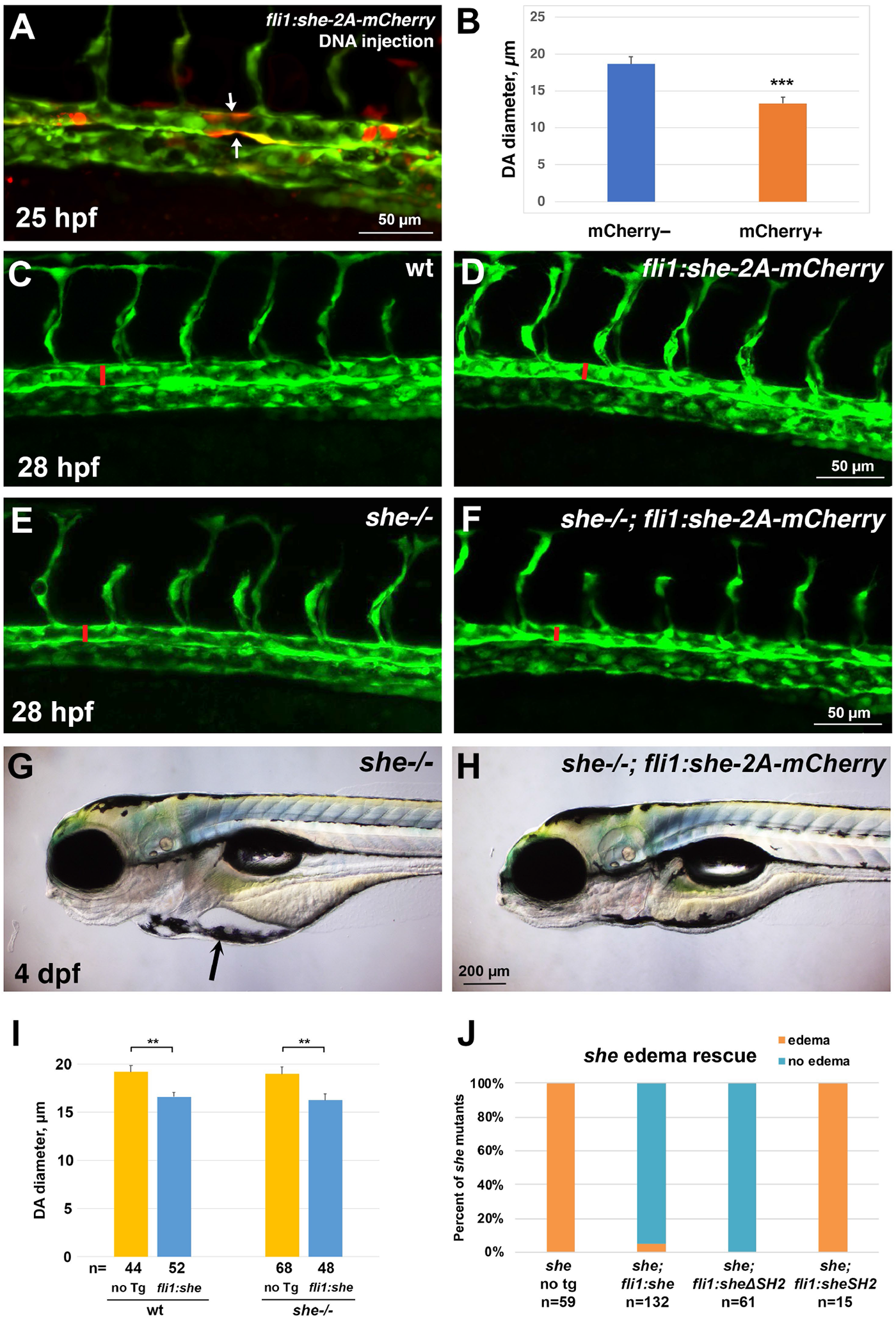
*she* overexpression in vasculature reduces the diameter of the dorsal aorta and rescues the *she* mutant phenotype. (A) Injection of *fli1:she-2A-mCherry* DNA plasmid together with Tol2 mRNA results in mosaic expression of *she-2A-mCherry* in endothelial cells. Note that mCherry positive segments of the DA (arrows, A) show reduced diameter compared to adjacent mCherry-negative segments. (B) Quantification of the DA diameter in mCherry+ and mCherry– DA segments in embryos from two independent experiments. n denotes the number of embryos analyzed. *** p<0.001 (Student’s t-test). (D-F) Trunk vasculature of wild-type*, fli1:she-2A-mCherry, she-/-* and *she-/-; fli1:she-2A-mCherry* embryos in *kdrl:GFP* background at 28 hpf. Confocal imaging of GFP expression in live embryos obtained from stable transgenic lines; the diameter of the DA is marked with red line. (G,H) Pericardial edema in *she-/-* mutants (E) is rescued in *she-/-; fli1:she-2A-mCherry* embryos at 4 dpf. (I) DA diameter is reduced in *fli1:she-2A-mCherry* embryos compared to wild-type non-transgenic (no Tg) siblings. Similarly, DA diameter is reduced in *she-/-; fli1:she-2A-mCherry* embryos compared to *she* siblings without the transgene. Embryos were analyzed at 28 hpf in *kdrl:GFP* background. Embryos were obtained by incross of either *she+/+* (referred to as wild-type); *fli1:she-2A-mCherry+/-* or *she-/-; fli1:she-2A-mCherry+/-* parents and separated based on mCherry expression. Note that wild-type and *she-/-* embryos cannot be directly compared in this experiment because they came from different parents and are not siblings; embryos from different pairs can show significant variability in the DA diameter. Error bars show SEM. **p<0.01, Student’s t-test. (J) Percentage of *she* mutant embryos showing pericardial edema at 4 dpf. Note that vascular endothelial expression of full length She (*fli1:she-2A-mCherry)* and the construct carrying a deletion of the SH2 domain in She protein *(fli1:she1′SH2-2A-mCherry)* rescues the mutant phenotype, while overexpression of She construct with SH2 domain alone *(fli1:sheSH2-2A-mCherry)* fails to rescue the phenotype.

SH2 domains are known to bind to phosphorylated tyrosines [26]. To test the functional requirement for the SH2 domain within the zebrafish She protein, we created She deletion constructs (Suppl. Fig. S4D) and generated stable zebrafish transgenic lines using vascular endothelial *fli1* promoter. Intriguingly, the construct that lacked the SH2 domain was sufficient to rescue the *she* mutant phenotype, and no pericardial edema or circulation defects were observed in *fli1:sheΔSH2-2A-mCherry; she-/-* embryos, which were viable through adulthood (Fig. 3H). In contrast, the construct containing only the SH2 domain was insufficient to rescue the *she* mutant phenotype (Fig. 3H).

To determine when She function is required, we made a construct which encodes a full-length zebrafish She under a heat-shock inducible *hsp70* promoter followed by 2A-mCherry and established stable zebrafish lines using Tol2 transgenesis. Heat-shock at 3 or 3.5 dpf was sufficient to rescue the pericardial edema phenotype fully or partially in *she* mutant embryos, positive for the *hsp70:she-2A-mCherry* transgene (Fig. 4A,B,G). This also prevented the collapse of DA at 4 dpf (Fig. 4C-G). This suggests that She has a critical functional requirement between 3 and 4 dpf.

**Figure 4.**
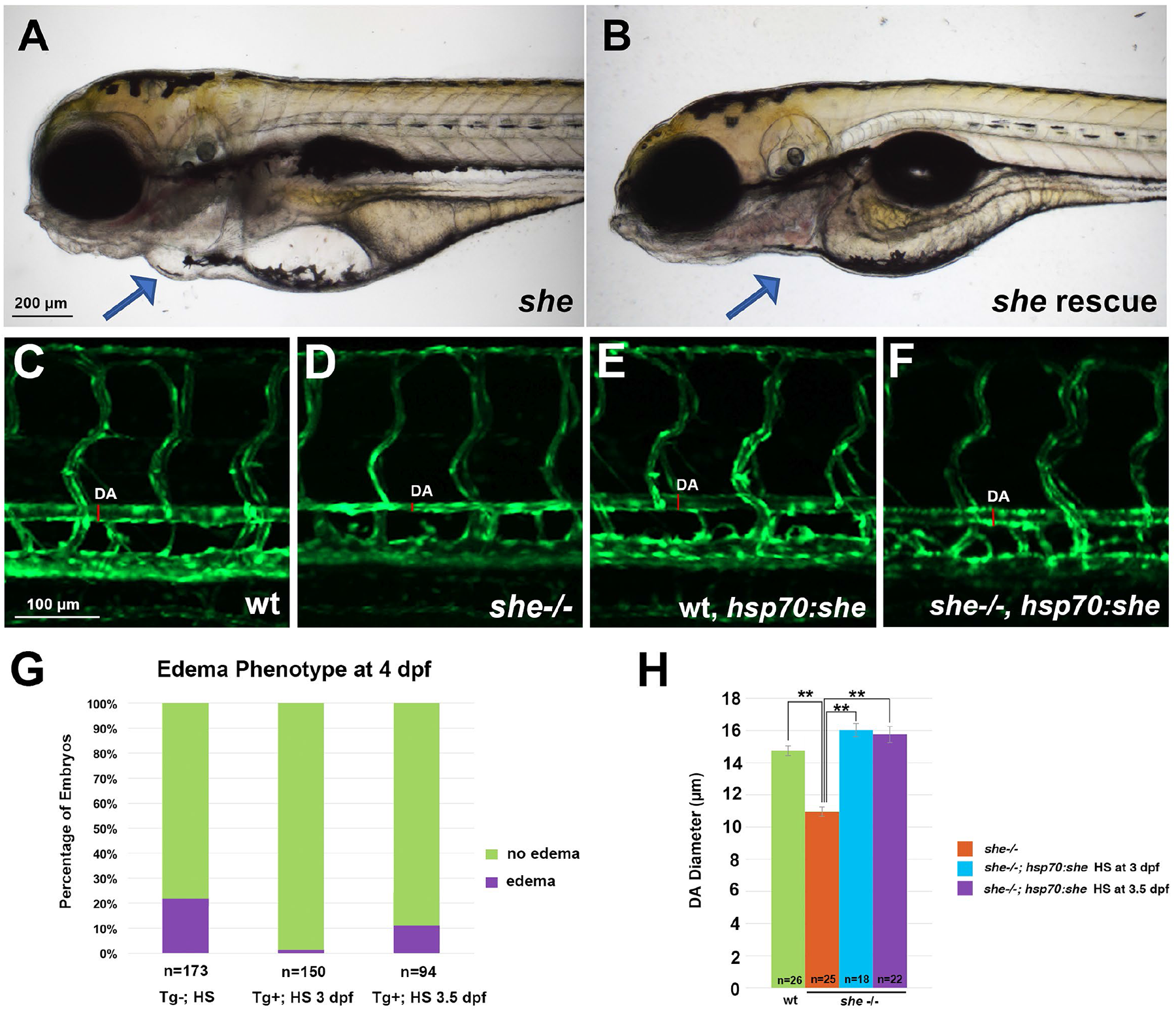
Inducible She expression under a heat-shock promoter *hsp70* rescues the *she* **mutant phenotype.** (A,B) Pericardial edema is observed in *she* mutant embryos at 4 dpf (A), while embryos positive for *hsp70:she-2A-mCherry* do not show pericardial edema and have normal blood circulation after heat-shock was performed at 3 dpf (B). Embryos were obtained from the cross of *hsp70:she-2A-mCherry; cryaa:dsRed +/-; she+/-* X *she+/-* parents in *kdrl:GFP* background. Embryos were sorted for dsRed in the lens (driven by the lens specific *cryaa* promoter which correlates with the presence of *hsp70:she-2A-mCherry* transgene) and mCherry in the body. Representative embryos with and without edema are shown for each group. (C-F) Dorsal aorta in *she* mutants is narrower or collapsed at 4 dpf, while heat-shock at 3 dpf restores normal DA size in *she-/-; hsp70:she-2A-mCherry* embryos. (G) Percentage of total embryos showing the pericardial edema phenotype at 4 dpf. Embryos were obtained by crossing *hsp70:she-2A-mCherry; cryaa:dsRed +/-; she+/-* X *she+/-* adults in *kdrl:GFP* background and subjected to heat-shock (HS) at 3 or 3.5 dpf. Embryos with dsRed in the lens (Tg+) or transgene negative controls (Tg-) were selected for the analysis. Tg-embryos were also subjected to heat shock at 3 or 3.5 dpf (both groups were combined). (H) The diameter of the dorsal aorta at 4 dpf in wild-type siblings (includes heterozygous and wild-type embryos), *she* mutant embryos without the transgene, and *she-/-; hsp70:she-2A-mCherry* embryos heat-shocked at 3 or 3.5 dpf. ** p<0.01, Student’s t-test.

To determine the cellular cause of the enlarged aorta observed in *she* mutants, we analyzed the endothelial cell number. The number of cells in the DA was significantly increased in *she* mutants crossed to the *kdrl:NLS-mCherry* endothelial nuclear reporter line when compared with their wild-type siblings at 72 hpf (Fig. 5A-C). In contrast, *fli1:she-2A-mCherry* embryos, which overexpress *she* in endothelial cells, had significantly fewer cells in the DA compared to their sibling mCherry negative embryos in the nuclear *fli1:NLS-GFP* reporter background (Fig. 5D-F). As analyzed by BrdU incorporation assay at 30 hpf*, fli1:she-2A-mCherry* embryos in *she* mutant background (which are phenotypically normal) showed reduced cell proliferation within the DA compared with mCherry-negative *she*-/-sibling embryos (Fig. 5G-M). These results argue that *she* restricts the DA size by inhibiting endothelial cell proliferation. We also analyzed mural cell formation by performing fluorescent ISH analysis for *pdgfrb* expression. However, no significant change *pdgfrb* expression was observed, suggesting that the changes in the DA size are not caused by defects in mural cell formation (Suppl. Fig. S5).

**Figure 5.**
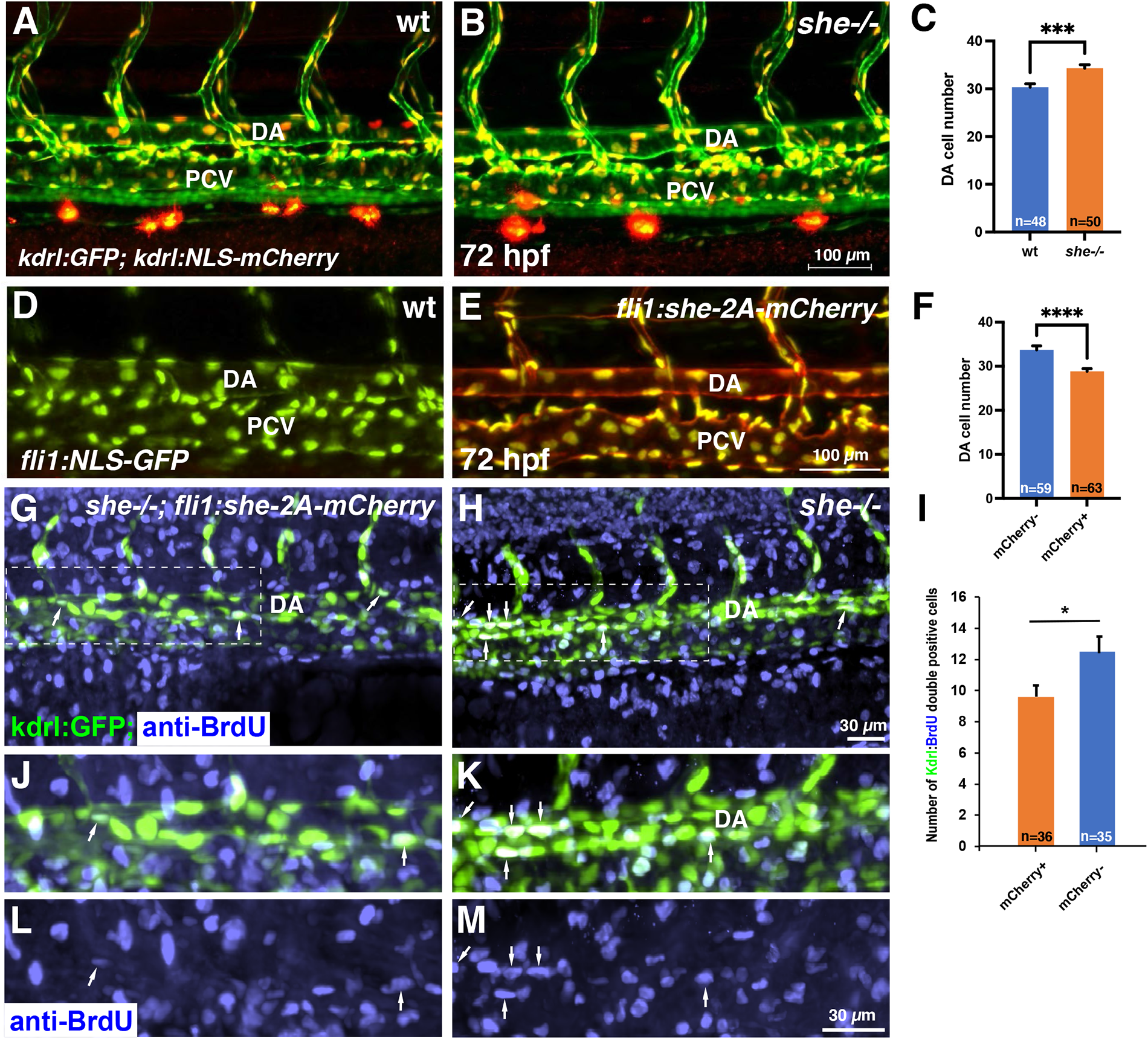
*she* affects cell number and inhibits endothelial cell proliferation. (A-C) Analysis of cell number in the DA of *she* mutant and wild-type *(she+/+)* sibling embryos at 72 hpf. Embryos were obtained from an incross of *she+/-; fli1:GFP; kdrl:NLS-mCherry* parents, imaged by confocal microscopy and subsequently genotyped. Note increased cell number in *she* mutant embryos. Data combined from 2 independent experiments. ***p<0.001, Student’s t-test. (D-F) Analysis of cell number in the DA of *fli1:she-2A-mCherry; fli1:NLS-GFP* embryos and their mCherry negative (wt) siblings at 72 hpf. Note the reduced cell number in mCherry+ embryos. Data combined from 2 independent experiments. ****p<0.0001, Student’s t-test. (G-M) Cell proliferation analysis using BrdU incorporation assay in *she-/-; fli1:she-2A-mCherry* embryos (phenotypically normal) and their sibling mCherry negative *she-/-* embryos in *kdrl:GFP* background. Note the increased number of BrdU and *kdrl:GFP* double positive cells within the DA in mCherry-negative *she* mutant embryos. Data combined from 2 independent experiments. *p<0.05, Student’s t-test.

It has been previously demonstrated that SHE interacts with ABL kinase in vitro [24]. To test a potential role for ABL signaling in regulating vascular lumen size, we treated zebrafish embryos with chemical inhibitors of ABL signaling, dasatinib and GNF-7 [27, 28], starting at 6 hpf. Embryos treated with either ABL inhibitor displayed reduction in the DA size at 28 hpf (Fig. 6A-E). No other morphological defects were observed at this stage. By 4 dpf dasatinib and GNF-7-treated embryos developed pericardial edema and their blood circulation slowed down or stopped, while the DA diameter remained reduced at this stage (Fig. 6F-H). Importantly, the number of endothelial cells in the DA was reduced in GNF7 treated embryos at 28 hpf, compared to DMSO-treated controls (Fig. 6I-K).

**Figure 6.**
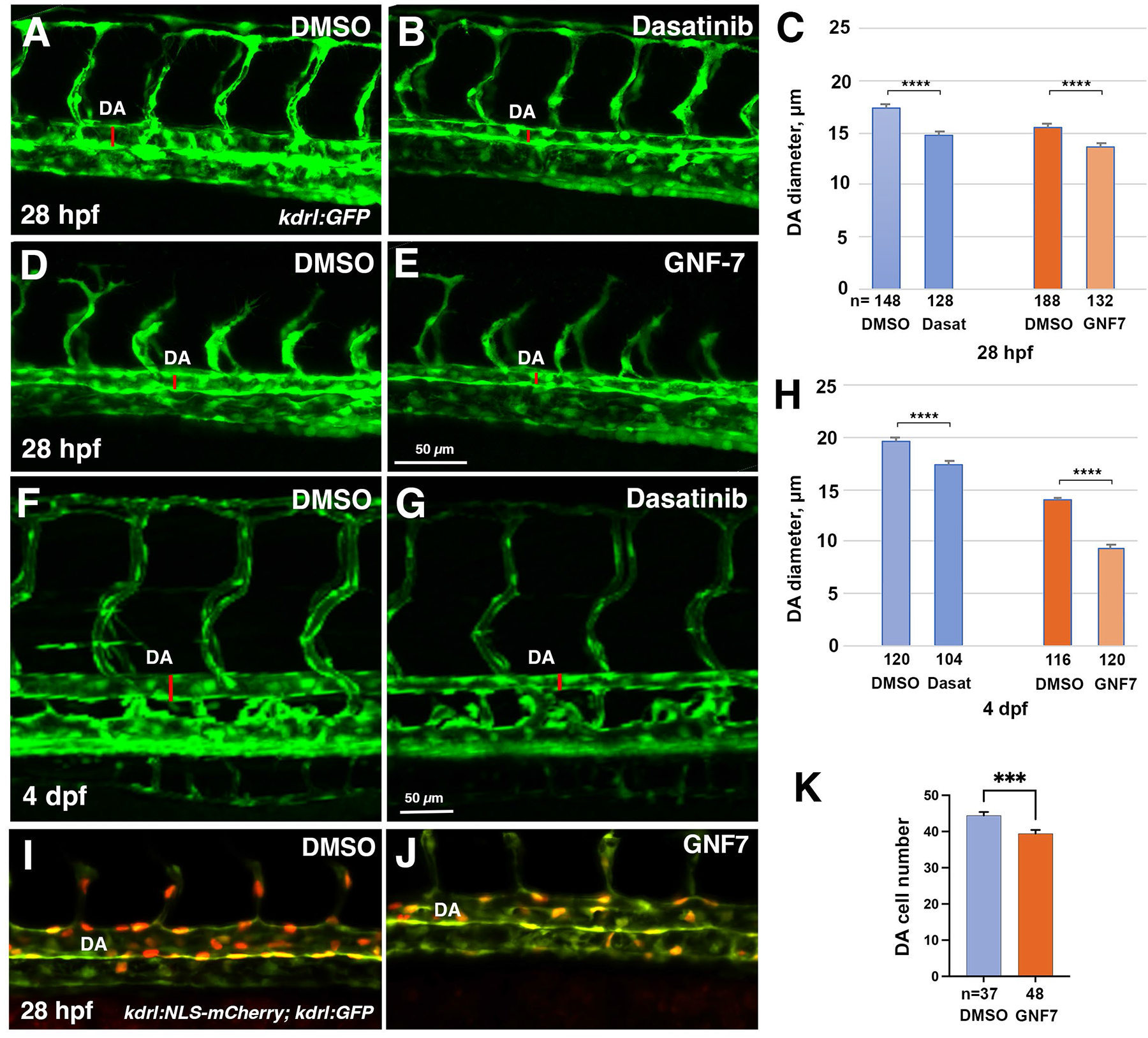
Inhibitors of Abl signaling reduce DA diameter and cell number in zebrafish embryos. (A-E) Dorsal aorta diameter at 28 hpf is reduced in *kdrl:GFP* embryos treated with 5 µM Dasatinib or 0.5 µM GNF-7 compared to controls treated with 0.1% DMSO. (F-H) Embryos treated with 5 µM Dasatinib exhibit pericardial edema (arrow) and narrower DA at 4 dpf compared to controls treated with 0.1% DMSO. (I-K) The number of cells in the DA is reduced in embryos treated with 1 µM GNF-7 compared to control embryos treated with 0.1% DMSO. Error bars show SEM in all graphs. ***p<0.001, ****p<0.0001, Student’s t-test. Data are combined from at least two independent experiments.

These results show that both She overexpression and inhibition of Abl signaling resulted in similar phenotypes of reduced DA diameter and lowered arterial cell number. We tested if inhibition of Abl signaling may reverse the enlarged DA observed in *she* mutants. *she* mutants and wild-type sibling embryos were treated with GNF-7 and analyzed for the DA diameter at 2 dpf. Indeed, while *she* mutants showed enlarged DA, GNF-7 treatment reversed this phenotype and resulted in a reduced width DA (Fig. 7).

**Figure 7.**
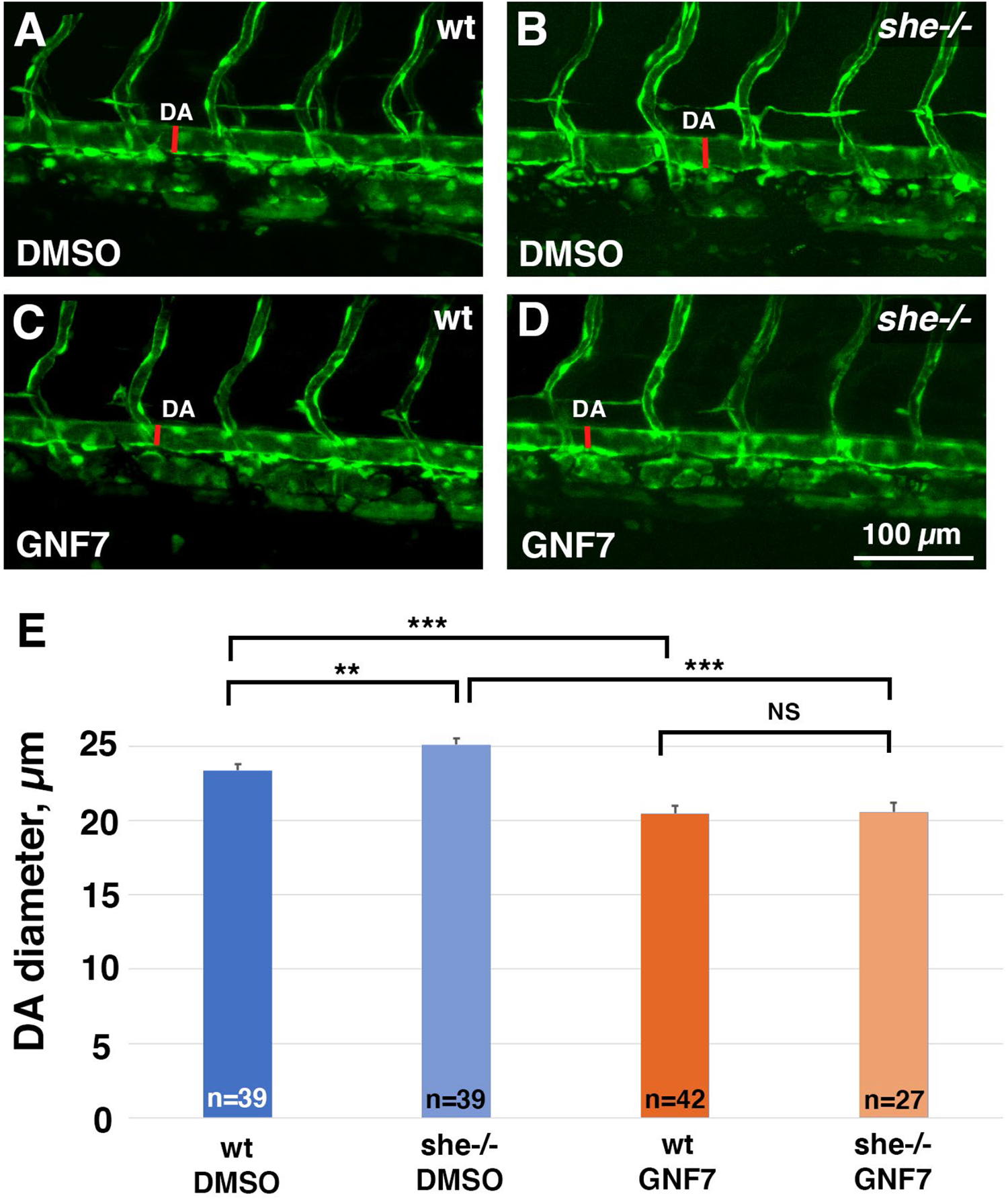
Inhibition of Abl signaling reverses enlarged DA in *she* mutant embryos. (A-D) *she+/-; kdrl:GFP* adults were crossed to obtain *she* mutant embryos. Embryos were treated starting at 6 hpf with either 0.5 µM GNF-7 or 0.1% DMSO. Embryos were imaged at approximately 55 hpf and subsequently genotyped. DA measurements were performed blinded. Mid-trunk region is shown, anterior is to the left. Note the slightly wider DA (red line) in *she-/-* mutant embryos compared to wild-type *(she+/+)* siblings. DA is reduced in both wild-type and *she* mutant embryos treated with GNF-7. (E) Quantification of DA diameter in wild-type or *she* mutant embryos treated with GNF-7 or DMSO. Error bars show SEM. **p<0.01, ***p<0.001, NS – not significant, Student’s t-test. Summary of two independent experiments.

It has been previously shown that ABL kinase can phosphorylate a related protein SHD in vitro [24]. The SHE sequence has four consensus tyrosine kinase phosphorylation sites with the YXXP motif (Suppl. Fig. S6A), which are conserved between different vertebrates and between SHE and SHD. Therefore, we hypothesized that ABL kinase may similarly target SHE protein for phosphorylation. To test the in vivo requirement for the consensus phosphorylation sites, we performed site directed mutagenesis on the zebrafish She to substitute all four consensus tyrosines into phenylalanine (YXXP –> FXXP). A zebrafish line was established that expressed the mutated She FXXP sequence under *fli1* promoter. Expression of the mutant construct failed to rescue pericardial edema and blood circulation defects in *she* mutant embryos (Suppl. Fig. S6B). In contrast to overexpression of wild-type *fli1:she-2A-mCherry* construct which results in narrower DA, *fli1:sheFXXP-2A-mCherry* embryos showed DA enlargement at 28 hpf stage (Suppl. Fig. S6C). Overexpression of the mutant SheFXXP may interfere with the wild-type She protein, resulting in a dominant negative effect, which can explain the DA enlargement observed in *fli1:sheFXXP* embryos. Altogether, these results argue that the consensus YXXP phosphorylation sites are required for SHE function.

Recently it has been reported that zebrafish mutants for claudin *cldn5a*, the homolog of the mammalian Cldn5, exhibit reduced DA [29]. Therefore, we tested if *cldn5a* expression was altered in *she* mutants. Indeed*, she* mutants exhibited increased *cldn5a* expression within the DA at 24 hpf when analyzed by standard chromogenic and fluorescent hybridization chain reaction (HCR) in situ hybridization approaches (Fig. 8A-D). Average *cldn5a* fluorescence intensity was increased by 17% in *she* mutants compared to wild-type embryos (Fig. 8N). Expression of *cldn5a* in the neural tube was unaffected in *she* mutants. To verify this result, we purified all vascular endothelial cells using FACS sorting from *kdrl:GFP* line in *she* mutants and wild-type embryos at 24 hpf and performed qPCR. 1.4-fold increase in *cldn5a* expression was observed using this approach (Fig. 8M). Arterial expression of a related homolog *cldn5b* was also increased in *she* mutants (Suppl. Fig. S7), and so was Cldn5 protein expression (both Cldn5a and Cldn5b are recognized by the same Cldn5 antibody) (Suppl. Fig. S8). In contrast, inhibition of Abl signaling by treating wild-type *kdrl:GFP* embryos with GNF7 resulted in a reduced *cldn5a* expression in the DA (Fig. 8E-H, O). To test if reduction in *cldn5a* level could restore the DA size, we knocked down *cldn5a* function in *she* mutants and sibling embryos using a previously validated Cldn5a morpholino (MO) [29]. A subphenotypic dose of MO was used which did not result in a significant reduction of DA width in wild-type embryos. Intriguingly, the same dose of *cldn5a* MO resulted in a reduction of DA diameter in *she* mutants leading to a partial rescue of the *she* phenotype (Fig. 8I-L, P).

**Figure 8.**
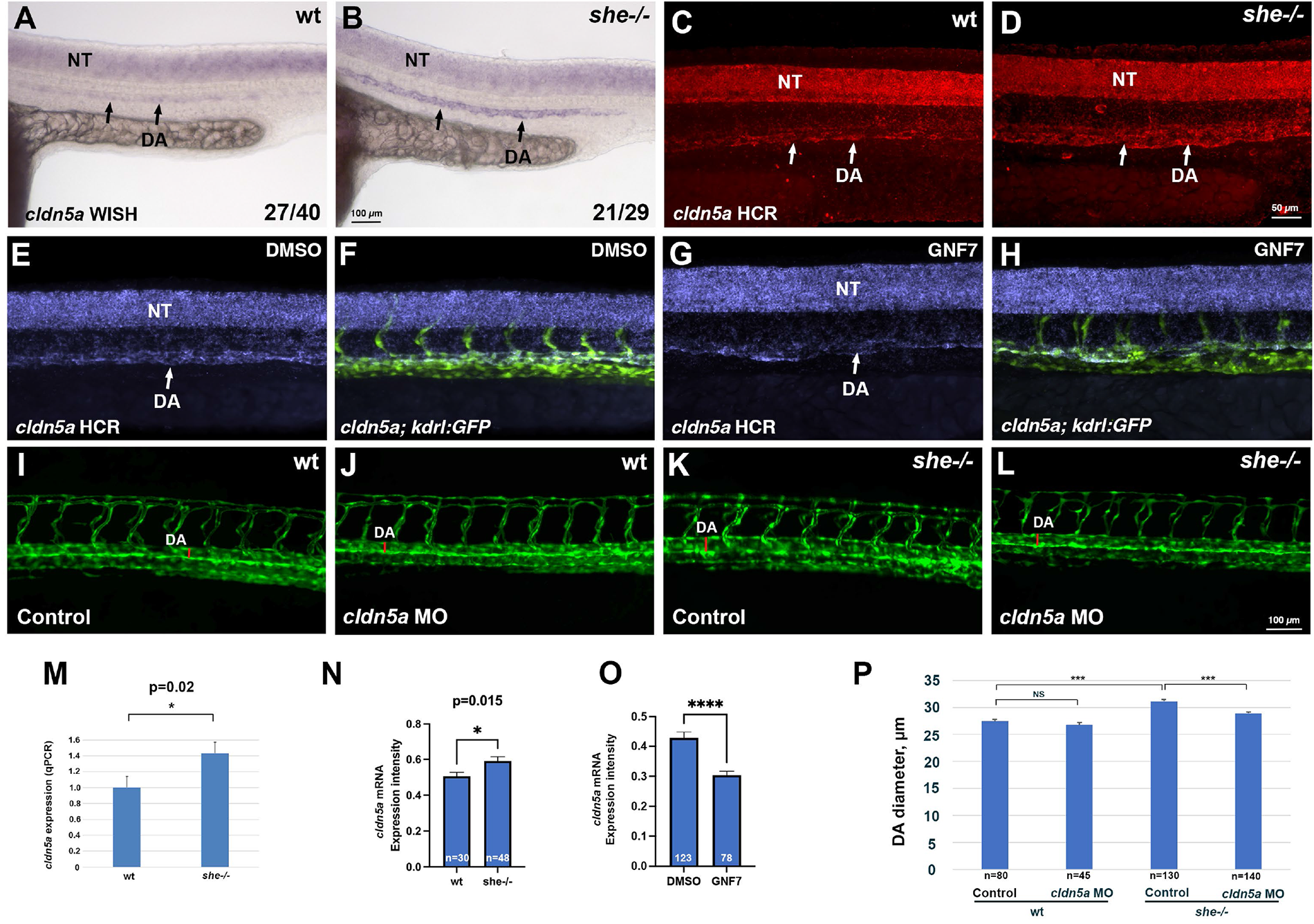
She regulates vascular lumen size by inhibiting *cldn5a* expression. (A-D) Chromogenic whole mount in situ hybridization (WISH) (A,B) and fluorescent in situ hybridization analysis using hybridization chain reaction (HCR) (C,D) analysis for *cldn5a* expression in *she* mutants and sibling wild-type *(she+/+)* embryos at 24 hpf. Note increased *cldn5a* expression in the dorsal aorta (DA) in *she* mutants while neural tube (NT) expression is unaffected. Quantification of *cldn5a* expression in the DA is shown in (N). Embryos were obtained by incross of heterozygous *she* parents and genotyped after WISH or HCR. (E-H) HCR analysis for *cldn5a* expression in control 0.01% DMSO (E,F) or 1 µM GNF7 treated *kdrl:GFP* embryos (G,H). *cldn5a* (purple) and merged *cldn5a; kdrl:GFP* channels are shown in the trunk region, anterior is to the left. Note the reduction in *cldn5a* expression in the DA while neural tube expression is unaffected. Quantification is shown in (O). (I-L) Inhibition of Cldn5a function using *cldn5a* MO injection reduces enlarged DA in *she* mutants. Embryos were obtained by an incross of *she+/-; kdrl:GFP* parents and imaged for GFP expression at 2 dpf. Mid-trunk region is shown, anterior is to the left. Quantification is shown in (P). Note the increase in DA width (red line) in *she* mutants (K), which is reduced upon *cldn5a* MO injection (L). (M) qPCR analysis of *cldn5a* expression in vascular endothelial cells obtained from FACS sorted *she* mutant and wild-type embryos at 24 hpf. Embryos were obtained by incross of *she-/-; fli1:she-2A-mCherry+/-* or *she+/+* (wild-type); *fli1:she-2A-mCherry+/-* parents in *kdrl:GFP* background and sorted for the absence of mCherry. Error bars show SEM. (N) Quantification of *cldn5a* expression in the DA of *she* mutants and wild-type siblings after HCR fluorescent in situ hybridization at 24 hpf. To reduce the staining variability between different embryos, expression values in the DA were normalized to the neural tube expression in each embryo. Expression analysis was performed blindly without knowledge of embryo genotypes. (O) Quantification of *cldn5a* expression in embryos treated with 0.01% DMSO or 1 µM GNF7 at 24 hpf. Relative expression in the DA was calculated by dividing the intensity in the DA over the expression intensity in the neural tube. Student’s t-test, error bars show SEM in (N,O). (P) Quantification of DA diameter at 2 dpf in wild-type and *she* mutant embryos injected with *cldn5a* MO. Error bars show SEM. *** p<0.001, NS – not significant.

To test the conservancy of She function in mammalian cells, we inhibited She function in human umbilical vein endothelial cells (HUVECs) using siRNA. Cells were transfected with either She or control siRNA, and tubulogenesis was assayed using 3D collagen matrix assay under serum-free defined conditions. siRNA transfected cells showed enlarged vascular lumens compared to control cells, reproducibly observed both at 48 and 72 hours after transfection (Fig. 9A-F). Assay of intracellular signaling kinases by Western blot showed increased phosphorylation of pPAK4 (Fig. 9G,H), which is a known signaling effector acting downstream of Cdc42 during vascular tubulogenesis [7, 8, 30]. Importantly, phosphorylation of a known ABL signaling effector CRKL was also increased in SHE inhibited cells, while pERK phosphorylation was not affected (Fig. 9G,H). These results argue that She function is conserved between vertebrates and suggest that She restricts lumen size by inhibiting Abl and Cdc42 activities.

**Figure 9.**
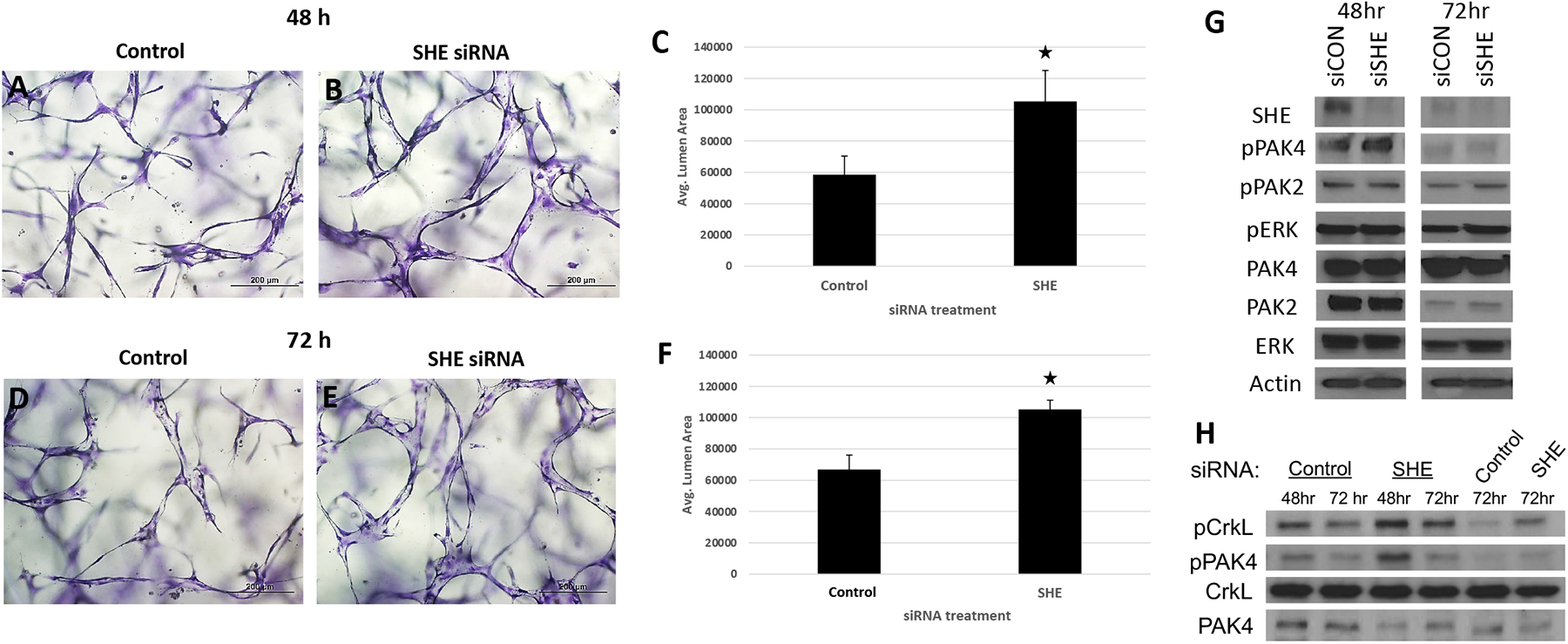
Inhibition of SHE in HUVECs results in enlarged tubulogenesis. (A-F) HUVEC cells were transfected with either control or SHE siRNA and analyzed in 3D collagen matrix assay at 48 (A-C) or 72 h (D-F) after transfection. The values (± standard deviation) are derived from analyzed fields (n=15) where total EC tube area was measured. Asterisks indicate significance at p<0.05 compared to control conditions. (G,H) Western blots for expression of SHE and phosphorylation of PAK4, PAK2, ERK and CRKL in HUVEC cells transfected with a control or SHE siRNA. Note greatly reduced SHE expression and increased pPAK4 and pCRKL in cells transfected with siSHE RNA. (H) shows replicate experiments for pPAK4 and pCRKL.

## Discussion

Our results demonstrate that She functions as a negative regulator of vascular lumen size. Loss of function *she* zebrafish mutants exhibit enlarged DA and have more arterial cells, while vascular endothelial *she* overexpression resulted in a reduced size DA with fewer cells. Inhibition of Abl signaling resulted in a similar reduction in DA size and cell number in wild-type embryos and reversed DA enlargement observed in *she* mutants, suggesting that She functions as a negative regulator of Abl signaling. A previous study demonstrated a direct interaction between She and Abl kinase in a yeast two hybrid assay [24]. Abl binding partners are often also substrates and/or activators of Abl kinase. Based on our data, we hypothesize that *she* functions as an adaptor protein to inhibit Abl activity. According to this model, Abl phosphorylates She, which then acts as a negative regulator to reduce Abl activity (Fig. 10). This type of feedback inhibition would allow for more accurate and sensitive regulation of vascular lumen size and could be modulated by She expression or its phosphorylation level.

**Figure 10.**
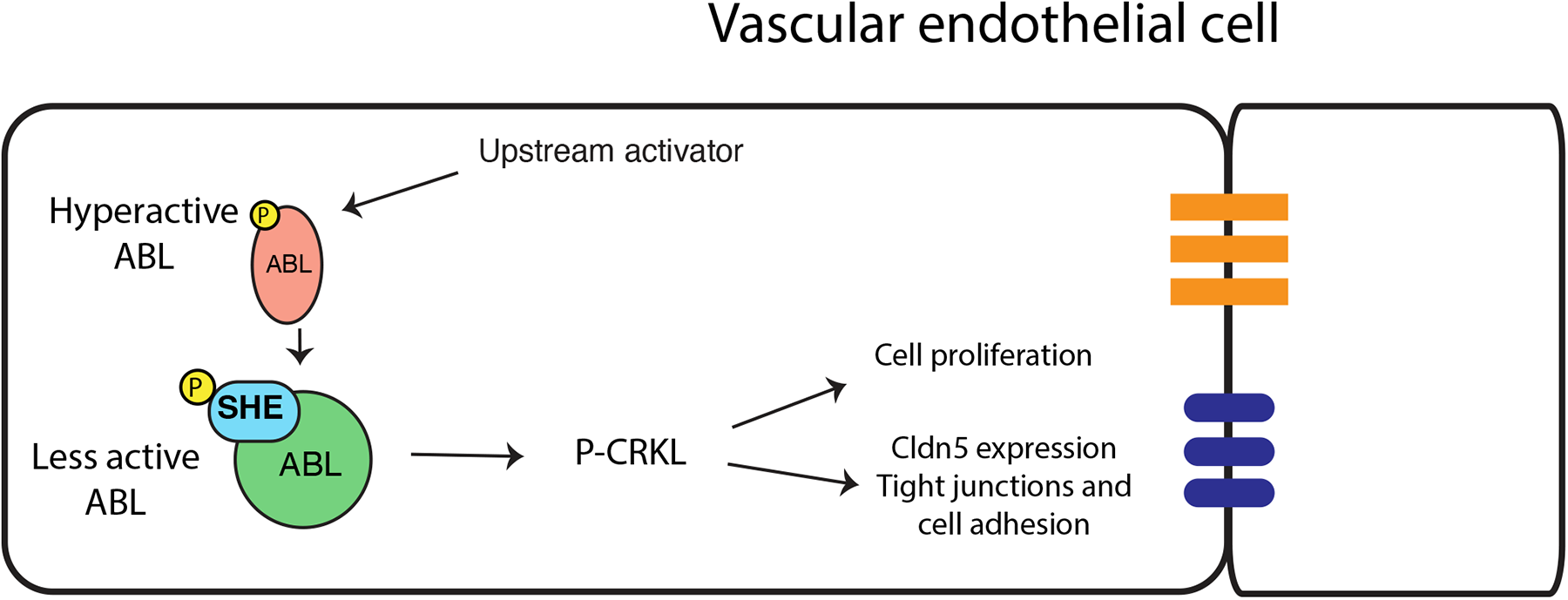
A proposed model for SHE and ABL signaling during vascular tubulogenesis. Activated ABL promotes enlarged vascular lumen through a downstream effector P-CRKL which increases endothelial cell proliferation and increases Cldn5 expression, thus affecting tight junctions and cell adhesion. Activated ABL phosphorylates SHE, which then interacts with ABL to dampen its activity resulting in the lumen of appropriate size.

In support of this model, zebrafish She has four YXXP motifs that correspond to predicted Abl tyrosine phosphorylation sites [31], all of which are conserved in human and mouse. She mutant variant YXXP -> FXXP with substitutions of tyrosines to phenylalanines failed to rescue the *she* mutant phenotype, demonstrating the importance of all four consensus phosphorylation sites for She function. Notably, these ABL phosphorylation sites are located outside of the SH2 domain, which is consistent with our results showing that the She protein lacking the SH2 domain is sufficient to rescue *she* mutant embryos. These data strongly suggest that Abl kinase phosphorylates She on these consensus sites, which is then critical for She activity.

Although the role of Abl signaling in normal vascular tubulogenesis has not been previously known, recent work has established the role of ABL signaling during pathogenesis of venous malformations (VM). In a VM cell culture model which overexpresses activating Tie2 receptor with Tie2-L914F mutation, enlarged vascular tubes and increased phosphorylation of c-ABL was observed [23]. Inhibition of ABL signaling reduced vascular lumen size and caused VM regression. Our data suggest that ABL signaling may play a similar role during embryonic tubulogenesis by promoting enlargement of vascular lumen, while the upstream initiating signal remains unclear.

Abl kinase is known to signal through multiple cytoplasmic effector proteins in different cellular contexts. In endothelial cells, Abl can enhance Angiopoietin-mediated activation of the AKT and ERK signaling pathways [20]. It is also known to promote Rac1 activation and cell-cell adhesion by phosphorylating Crk and Crkl proteins, which are considered specific substrates of Abl kinase [15, 32, 33]. Our results show that Crkl phosphorylation is increased in She knockdown HUVEC cells, which suggests that Crkl functions as a downstream effector of She-Abl signaling in vascular tubulogenesis.

While our results strongly suggest that She functions in the Abl signaling pathway, it is important to note that the chemical inhibitors may also inhibit additional signaling pathways, and their specificity in vivo has not been fully determined. Therefore, we cannot exclude the possibility that the DA decrease in GNF7 and dasatinib treated embryos may be caused by inhibition of other signaling pathways. Obtaining additional genetic evidence will be important to fully validate the role of Abl signaling in vascular tubulogenesis.

Rho GTPases (Rac1, Cdc42 and RhoA) have been previously implicated as key regulators of actomyosin network during vascular lumen formation [1–5]. Rac1 and Cdc42 stimulate cytoskeletal remodeling to establish cell polarity including localization and stabilization of cell junctions to regulate endothelial tubulogenesis, while RhoA has an opposing activity [3, 8, 34]. Abl signaling has been implicated in regulating Rac1 activity in different non-endothelial cell types [13–16], therefore it is likely that She and Abl signaling regulate vascular lumen size by modulating the activity of Cdc42 and Rac1 or other GTPases. In support of this model, our data demonstrate that phosphorylation of PAK4, a downstream effector of Cdc42 [7, 30], is increased in *SHE* knockdown endothelial cells.

In addition to regulating cytoskeletal remodeling, Abl signaling is known to promote cell proliferation in different cell types, including smooth muscle cells and fibroblasts [18, 19]. Our data argue that Abl signaling also promotes vascular endothelial cell proliferation in the DA, while She acts as an inhibitor of this process. It is quite likely that downstream effectors affecting cell proliferation and cytoskeletal rearrangements are different, which needs to be investigated further in the context of vascular tubulogenesis.

Intriguingly, *she* expression is primarily enriched in the arterial vasculature, including the DA [25]. *she* mutants had little if any change in the PCV diameter, while DA diameter was enlarged as early as 28 hpf. Abl inhibition did not cause a significant change in the PCV diameter, while the DA was reduced. This suggests that Abl activity primarily regulates size of the arterial vasculature. Although vascular diameter in older embryos and adults is regulated by smooth muscle contraction, there is no smooth muscle cell coverage yet at 28 hpf. Therefore, DA enlargement is unlikely to be caused by defects in smooth muscle cells. Our results show that *cldn5a* and *cldn5b* expression, which encode zebrafish homologs of Claudin 5, a major component of the tight junctions, is upregulated in *she* mutants. It is possible that She-Abl signaling directly represses *cldn5a/b* expression to ensure an optimal amount of Cldn5a/b. Alternatively, *cldn5a/b* expression may be upregulated as a secondary consequence of other changes in cell shape, junctions, or polarity. *cldn5a* mutants display reduction in the DA lumen size [29]. Our results show that downregulation of *cldn5a* expression reverses the phenotype of enlarged DA observed in *she* mutant embryos, suggesting the DA enlargement is mediated at least partially by an increase in *cldn5a* expression. In addition to mediating vascular permeability, Cldn5 has been implicated in cell motility and proliferation of endothelial and tumor cells [29, 35, 36]. It is possible that increased cell proliferation in the DA may be a direct or indirect consequence of upregulated Cldn5 expression. Alternatively, increase in Cldn5 expression may be independent from She effect on cell proliferation.

The SHE protein sequence is highly conserved between zebrafish and other vertebrates. Intriguingly, siRNA knockdown of human SHE in HUVECs resulted in a similar increase in vascular tube diameter. There is no blood flow or smooth muscle / pericyte cells in this 3D collagen assay, therefore the observed tube enlargement must be cell-autonomous to vascular endothelial cells. Endothelial-specific expression of She rescued the *she* loss-of-function phenotype in zebrafish embryos, further supporting cell autonomous role for SHE.

In summary, our results demonstrate novel roles for SHE and ABL signaling in regulating vascular lumen size. These findings will contribute to our understanding of molecular mechanisms that regulate vascular tubulogenesis and eventually may facilitate development of novel therapeutic approaches for pathological disorders, such as venous malformations which display ectatic vessels with enlarged lumens.

## Materials and Methods

### Zebrafish lines. Generation of she^ci26^ line

All zebrafish work was performed after approval of USF and CCHMC IACUC committees. TALEN genome editing was used to create the *she^ci26^* allele. TALENs were designed using the TAL Effector Nucleotide Targeter software (https://tale-nt.cac.cornell.edu/) [37]. The TALEN target sequence TGTGCGTATTCGTCATGGCgaagtggtttaaagaTTTTCCAACTAGTTTGAAGA is located in exon 1 of the *she* gene. The lowercase letters represent the spacer sequence, which contains MseI restriction site TTAA. TALENs were assembled using Golden Gate assembly [38], and mRNA for the left and right arms was synthesized using the T3 mMessage mMachine kit (ThermoFisher). 100 pg of each mRNA was injected into one-cell stage wild type embryos. she_F1 and she_R1 primers were used to test TALEN efficacy and identify *she^ci26^* adult carriers (primer sequences listed in Suppl. Table S1). These primers amplify a 333 bp product, which can then be digested with MseI.

### Generation of she^ci30^ line

CRISPR/Cas9 editing was used to create the *she^ci30^* allele. Guide RNAs were designed using ChopChop (http://chopchop.cbu.uib.no). Two separate guide RNA templates were generated by PCR as described [39], using she_gRNA1, she_gRNA2 and the common she_gRNA_R1 primer (Suppl. Table S1). gRNAs were synthesized using the MEGAshortscript T7 kit (Life Technologies). The two gRNAs were injected together with Cas9-NLS protein (New England Biolabs) into one-cell stage embryos. The injected animals were outcrossed and PCR was performed on pooled embryos to identify carriers of *she* mutations, using She_gRNA left and She_gRNA right primers (Suppl. Table S1). These primers span the large deletion in *she^ci30^,* and thus generate an approximately 200 bp PCR product only from the mutant allele. To identify the wild type allele, a separate PCR reaction was performed using She_gRNA midR reverse primer (Suppl. Table S1). This primer is located within the deleted region, thus will amplify a 234 bp product only from the wild type allele.

### Generation of fli1:she-2A-mCherry and hsp70:she-2A-mCherry lines and DNA overexpression analysis

To create full length over-expression constructs, we amplified the *she* full length coding sequence from wild type cDNA using she_attB_F1 and she_attB_R3 primers (Suppl. Table S1) for Gateway cloning (Invitrogen). The PCR product was recombined via BP reaction into pDONR221. To make the *fli1:she-2A-mCherry* construct, the resulting middle entry vector (pDONR221-she) was then recombined via LR reaction with p5E-fli1ep and p3E-2AmCherry into the destination vector pDestTol2pA2 [40]. To make the hsp70-she-2A-mCherry construct, pDONR221-she, p5E-hsp70, and p3E-2A-mCherry were recombined via LR reaction into the destination vector pDestTol2pA; cryaa:dsRed. Induction of the *hsp70:she-2A-mCherry* transgene was confirmed by ubiquitous mCherry fluorescence, while expression of fli1:she-2A-mCherry was observed in vascular endothelial cells (Suppl. Fig. S4).

To analyze mosaic overexpression, embryos were injected with 30 ng of *fli1:she-2A-mCherry* DNA construct and 225 ng of Tol2 mRNA mixture. Embryos with endothelial mCherry expression were chosen for confocal imaging and further analysis. mCherry positive and negative segments of the DA (1 segment per embryo) were chosen for measurements in Fiji software. 3 measurements were performed for each segment analyzed, and measurements were then averaged to obtain a single data point used for further calculations.

To create She deletion constructs, we amplified portions of *she* from wild type cDNA. To generate she1′SH2 which contains SH2 domain deletion, she_attB_F1 and she_deltaSH2_attB_R primers were used. To generate sheSH2 construct which contains SH2 domain only, sheSH2_attB_F and she_attB_R3 primers were used (Suppl. Table S1). Each product was recombined via BP reaction into pDONR221. The resulting middle entry vectors were each then recombined via LR reaction with p5E-fli1ep and p3E-2AmCherry into the destination vector pDestTol2pA2. All constructs were confirmed by sequencing. All final constructs were injected with Tol2 mRNA into one-cell stage embryos derived from a cross of *she^ci26^* heterozygous animals. Injected embryos were raised to adulthood and outcrossed to produce embryos to screen for germline integration. Transgenic lines were identified based on vascular endothelial mCherry expression.

To create the she-FXXP construct, we performed sequential site-directed mutagenesis on each YXXP site starting with the pDONR221-she construct, using the QuikChange II Site-directed Mutagenesis Kit (Agilent). Each construct was fully sequenced before proceeding with the next round of mutagenesis. The primer pairs sheYXXP1-P4 used for mutagenesis of each site are listed in Suppl. Table S1. This construct was then combined via LR reaction with the p5E-fli1ep and p3E-2A-mCherry into the pDestTol2pA2 destination vector, then injected as described above. All *she* lines were maintained in *kdrl:GFP^s843^* background [41].

### Other lines

The following lines were used in the study: *kdrl:GFP^s843^* [41], *Tg(kdrl:NLS-mCherry)^is4Tg^* [42], *Tg(fli1:nGFP)^y7Tg^* [43].

#### Heat-shock induction

Embryos at the indicated stages were placed in 96 well PCR plates in embryo media with 5-10 embryos per well. To induce transgene expression, embryos were incubated at 37°C for one hour in a thermocycler. Following heat shock, embryos were transferred from PCR tubes to fresh media in petri dishes and sorted for dsRed expression in the lens or mCherry fluorescence in the body to indicate presence of the transgene. For DA diameter analysis, embryos were genotyped following confocal imaging and analysis.

#### Abl inhibitor treatments

Dasatinib (Selleckchem) and GNF-7 (Selleckchem) were resuspended in DMSO to a concentration of 10 mM. Stocks were further diluted into embryo water to the appropriate working concentrations as noted. DMSO was diluted 1:1,000 or 1:10,000 in embryo water as a control. Pools of 20 dechorionated *kdrl:GFP* or *she*-/-; *kdrl:GFP^s843^* embryos were placed in 1 mL diluted inhibitor or diluted DMSO in glass vials starting at 6 hpf. Embryos were placed on a nutator in the dark until 24-28 hpf, when they were imaged for analysis.

#### In situ hybridization and hybridization chain reaction analysis

Chromogenic whole mount in situ hybridization was performed as described [44]. *she* DIG-labeled antisense mRNA probe was synthesized as previously described [45]. To make ISH probe for *cldn5a*, 897 bp fragment corresponding to the coding sequence of *cldn5a* was amplified by PCR from zebrafish 24 hpf embryonic cDNA. The PCR product was subcloned into pCRII-TOPO using TOPO TA cloning kit (ThermoFisher). The construct was linearized using XhoI restriction endonuclease (ThermoFisher), and antisense DIG-labeled RNA was synthesized using SP6 RNA polymerase (Promega).

Fluorescent in situ hybridization using hybridization chain reaction (HCR) for *cldn5a* was performed as previously described [46, 47]. Embryos were obtained from an incross of *she+/-; kdrl:GFP* parents, imaged using 10x or 20x objective on a Nikon A1R or Nikon Eclipse confocal microscope and subsequently genotyped. To measure fluorescence intensity of *cldn5a* staining, a small rectangular area which included a portion of the DA was selected using Fiji (Image J). Three measurements at different locations of the DA (using the same size rectangular) in the trunk region were performed for each embryo. In addition, a similar size area was selected in the *cldn5a* expression domain within the neural tube (for normalization) and in the embryo region which did not show any specific *cldn5a* expression (for background subtraction). Integrated density was calculated for each area. Subsequently, background subtraction was performed by subtracting integrated density values obtained for the area which did not show specific *cldn5a* expression. All DA expression values were then normalized by dividing them over the neural tube expression of the same embryo. This procedure reduced variation due to staining intensity variability between different embryos. *cldn5a* expression in the neural tube was not expected to be affected, therefore it served as a reference point. Student’s t-test was used to calculate statistical significance between normalized integrated density values of *cldn5a* expression in the DA of wild-type and *she* mutant sibling embryos. All measurements were performed blindly without a prior knowledge of embryo genotypes.

A similar approach was used to measure *cldn5b* intensity. HCR probes and fluorescent 647nm hairpins were obtained from Molecular Instruments Inc (Los Angeles, CA). Because *cldn5b* only stains DA but not the neural tube, normalization was not performed in this case. All measurement values (3 per embryo at different locations of the DA) were used for statistical analysis of *cldn5a* and *cldn5b* expression.

HCR for *pdgfrb* expression in *she; kdrl:GFP* embryos was performed under a similar protocol. *pdgfrb* probe was obtained from Molecular Instruments and used with 594nm hairpins. Fiji / Image J was used to measure the mean intensity of *pdgfrb* expression within the DA region that spans approximately 4 ISVs in each embryo. Data were combined from 3 separate replicate experiments. Because the overall staining intensity varied between separate experiments, relative fluorescence intensity was normalized within each experiment. For this purpose, mean *pdgfrb* fluorescence intensity values were calculated for wild-type embryos within each experiment. Then the measured intensity values for each wild-type or mutant embryo were divided over the same mean wt intensity value calculated for each experiment. The resulting relative expression values were used for the graph and statistical calculations.

#### Cldn5 immunostaining

Cldn5 immunostaining was performed using a previously established protocol [48]. Embryos were obtained from a cross of *she+/-; kdrl:GFP* parents, imaged using confocal microscopy and subsequently genotyped. To measure fluorescence intensity of Cldn5 immunostaining, a small rectangular area which included a portion of the DA was selected using Fiji (Image J). 10 measurements at different locations of the DA (using the same size rectangular) in the trunk region were performed for each embryo. In addition, a similar size area was selected in the Cldn5 expression domain within the neural tube (for normalization) and in the embryo region which did not show any specific Cldn5 expression (for background subtraction). Mean fluorescence intensity was measured for each area. Subsequently, background subtraction was performed by subtracting mean fluorescence values obtained for the area which did not show specific Cldn5 expression. All DA expression values were then normalized by dividing them over the neural tube expression of the same embryo. This procedure reduced variation due to staining intensity variability between different embryos. Student’s t-test was used to calculate statistical significance between all normalized values of Cldn5 expression in the DA of wild-type / heterozygous (*she+/+* and *she+/-* embryos were combined into a single group) and *she-/-* sibling embryos. All measurements were performed blindly without prior knowledge of embryo genotypes.

#### Morpholino injection

4 ng of a previously validated *cldn5a* MO [29](GeneTools Inc, sequence AGGCCATCGCTTTCTTTTCCCACTC) was injected into embryos at 1-cell stage obtained from a cross of *she+/-; kdrl:GFP* parents. Embryos from injected and control uninjected groups were imaged at 2 dpf using confocal microscopy (Nikon Eclipse) and genotyped. DA lumen diameter was analyzed as described above. All analysis was performed blindly without knowing embryo genotypes.

#### qPCR analysis using FACS-sorted endothelial cells

Wild-type and *she* mutant embryos were obtained from respective incrosses of *she+/+; fli1:she-2A-mCherry; kdrl:GFP* and *she-/-; fli1:she-2A-mCherry; kdrl:GFP* parent fish (which were siblings). 15-20 mCherry-negative embryos were obtained for each group at 24 hpf stage. Single-cell suspension was obtained as previously described [49]. FACS sorting for GFP-positive cells was performed at the CCHMC FACS Core facility. Total RNA was purified using RNA purification kit (Norgen Biotek). cDNA was synthesized at CCHMC Gene Expression Core facility. Quantitative real-time PCR (qRT-PCR) was carried out using PowerUp SYBR Green Master Mix (Applied Biosystems) in a StepOnePlus Real-Time PCR System. Each reaction was repeated twice, and two biological replicates were used. qPCR amplification for *cldn5a* and EF1α was performed using the following primers: cldn5a-F (GACAACGTGAAAGCGCGGG), cldn5a-R (AGGAGCAGCAGAGTATGCTTCCC), EF1a-F (TCACCCTGGGAGTGAAACAGC) and EF1a-R (ACTTGCAGGCGATGTGAGCAG). Fold changes were calculated using a relative standard curve methods and normalized to EF1a expression.

#### Microscopy imaging

For brightfield imaging, *she^ci30^* and *she^ci26^* embryos were obtained by respective crosses of heterozygous parents. Live embryos were anesthetized in 0.016% Tricaine solution, mounted in 2% methylcellulose and imaged using Zeiss Axioimager (10x objective). Embryos were subsequently genotyped. For brightfield imaging of embryos after in situ hybridization, they were mounted in 0.6% low-melting-point agarose and imaged using 10x objective on a Nikon Eclipse compound microscope.

Fluorescent imaging of live embryos and fixed embryos after HCR was performed using Nikon confocal microscope (A1R or Eclipse). Embryos were mounted in 0.6% low melting-point-agarose. Nikon Elements AR Denoise.ai algorithm was used to remove noise from images with high background signal. Maximum intensity projections were obtained using Nikon Elements or Fiji (Image J) software. Image levels were adjusted using Adobe Photoshop 2021 to maximize brightness and contrast; in all cases similar adjustments in control and experimental embryos were performed.

#### Measurements of DA and PCV diameter

To analyze the DA and PCV diameter of *she^ci26^* mutants, embryos were obtained by a cross of heterozygous parents. Live embryos were anesthetized in Tricaine, mounted in 0.6% LMP agarose and imaged using Nikon A1 confocal microscope. Subsequently embryos were retrieved and genotyped. Measurements were performed in Fiji (Image J). Four measurements across the DA or PCV were taken in different areas of the mid-trunk for each embryo. Measurements were performed at the midpoint between two ISVs, every other ISV. Because the diameter of the DA (or PCV) can vary within each embryo, each measurement was treated as an independent datapoint for statistical calculations. Measurements were performed blindly without knowledge of embryo genotypes. DA diameter was measured similarly in embryos treated with Abl inhibitors. In the experiment where *she* mutant and control embryos were treated with GNF7 (Fig. 6), 3 measurements within the DA in each embryo were performed (embryos were imaged at higher magnification, and thus a smaller region of DA in the trunk was available for analysis). In the experiment where *she* and mutant embryos were injected with *cldn5a MO* (Fig. 7E-H), 5 measurements within the DA in each embryo were performed (lower magnification imaging was used for this analysis covering a larger region of an embryo).

#### Cell number analysis

To analyze arterial cell number in *she* mutants, embryos were obtained from an incross of *she+/-; kdrl:GFP; kdrl:NLS-mCherry* adults and imaged using confocal microscopy under a 20x objective at 72 hpf. Embryos were then extracted from the LMP agarose and genotyped. Nuclei cell counts analysis of the DA region was performed within a segment between 5 ISVs using Imaris 7.2 software. First, the region of interest was selected to limit the counts within the DA. An estimated diameter was calculated by measuring the size of selected cell nuclei within the Imaris. Then, the preliminary results were manually corrected to remove or add nuclei based on the confocal 3D image. Similar approach was used for cell counts in GNF7 treated embryos and DMSO controls using *kdrl:GFP; kdrl:NLS-mCherry* line.

To count arterial cells in She overexpressing embryos, double transgenic *fli1:she-2A-mCherry; fli1:NLS-GFP* embryos were obtained from a cross of separate transgenic parents. Confocal imaging at 72 hpf and cell counts were performed in GFP channel as described above.

#### Cell proliferation analysis

Embryos were obtained through an incross of *she-/-; fli1:she-2A-mCherry; kdrl:GFP* adult zebrafish. mCherry positive and negative embryos were segregated by screening under a fluorescent microscope. The embryos were treated with BrdU (Millipore-Sigma, Cat No. B5002) in a 24-well plate (20 embryos per well in a 200µl of 15 mM BrdU solution in 15% DMSO) from 29 to 29.5 hpf. At 29.5 hpf, the embryos were allowed to recover in embryo water for 30 minutes and fixed afterwards at 30hpf. The embryos were fixed in BT-fix for 2 hours at room temperature (RT). Fixed embryos were walked with gradual changes into 100% methanol and kept overnight at -20°C. The next day, embryos were walked into PBT (PBS + 0.1% Tween 20). The embryos were washed with PBT for 3X15 mins. This was followed by incubation in 2N HCL for 1 hour at RT. The embryos were washed with PBT for 3X5 mins. The embryos were then placed in blocking buffer for 2.5 hours. This was followed by overnight incubation in primary anti-BrdU antibody (ab6326, abcam, 1:250 dilution). The embryos were washed the next day with PBT for 6X15 mins. This was followed by overnight incubation in the goat anti-rat Alexa Fluor 647 secondary antibody (A21247, ThermoFisher, 1:500). The embryos were washed with PBT 4X15 min and kept at 4°C overnight. The embryos were imaged the next day on a Nikon Eclipse confocal microscope with 20X water immersion objective. The entire trunk region was imaged in separate segments (3 images / embryo). The double labelled cells were counted using the Imaris software 9.1 (Bitplane). A region of interest was drawn to encompass only the DA. Separate spots were created for *kdrl:GFP* positive cells and anti-BrdU labeled cells, respectively. Co-localization was defined as spots by no more than 4µm. Colocalization and cell identity was confirmed by manual analysis of the image. The data were obtained from two independent experiments.

#### Intersegmental vessel analysis

To count arterial and venous connections, *she^ci26+/^-; kdrl:GFP* adults were crossed to obtain a mixture of *she* homozygous, heterozygous and wild-type embryos, which were then mounted in 0.6% LMP agarose and imaged by confocal microscopy using a 10x objective at 72 hpf. Embryos were then extracted from the agarose and genotyped. Obtained images were analyzed blindly without knowing embryo genotype to determine arterial or venous connections of 10 consecutive pairs (20 ISVs per embryo.)

#### EC tube formation assays in 3D collagen matrices

Human umbilical vein endothelial cells (ECs) were grown and used from passages 3-6 as previously described [50]. Human ECs were seeded in 3D collagen matrices in the presence of a combination of recombinant growth factors, stem cell factor (SCF), interleukin-3 (IL-3), stromal-derived factor-1 alpha (SDF-1α), and fibroblast growth factor-2 (FGF-2), which were added to Medium 199 containing a 1:250 dilution of reduced serum supplement-2 [50], which has insulin as a component. The recombinant growth factors were obtained from R&D Systems and were added at the following concentrations: SCF, IL-3, and SDF-1α at 40 ng/ml and FGF-2 at 50 ng/ml. The details of the serum-free defined assay system set-up have been described previously [51, 52]. siRNA suppression of ECs was performed as described [50], using two rounds of transfection using either a control siRNA or an siRNA directed to She. Silencer select siRNAs (Invitrogen) were utilized for these experiments with a control siRNA (Negative control #2-catalog number 4390896) and a She siRNA (catalog number s43069). After siRNA treatment, ECs were seeded in 3D collagen matrices in the presence of the recombinant growth factors, and after 48 or 72 hr, cultures were either fixed with 3% glutaraldehyde in phosphate-buffered saline or collagen gels were plucked and lysates were prepared using SDS-PAGE Laemmli 1.5X sample buffer. Western blots were run and probed with antibodies directed to phospho-Pak2, phospho-Pak4, phospho-Erk, phospho-Crkl (Tyr207, Cat No. 3181S), total Pak2, total Pak4, total Erk and total Crkl (Cat No. 38710S) (Cell Signaling Technology). The anti-She antibody was from Novus and the anti-actin antibody was from Calbiochem. Fixed cultures were stained with 0.1% toluidine blue, and cultures were photographed using an Inverted Olympus microscope, and tube area was traced and quantitated using Metamorph software. Representative experiments were analyzed, and the data was obtained from triplicate wells while lumen area was measured from n=15 fields. Statistical analysis utilized a Student’s t-test and statistical significance was established with a p<0.05.

#### Reagent availability

All DNA plasmids and zebrafish lines are available from the corresponding author upon reasonable request.

## Acknowledgements

We thank Matt Kofron and CCHMC Confocal Core for assistance with imaging and CCHMC FACS core for assistance with cell sorting.

