## Supplemental Materials for "SH2 domain protein E (SHE) and ABL signaling regulate blood vessel size"

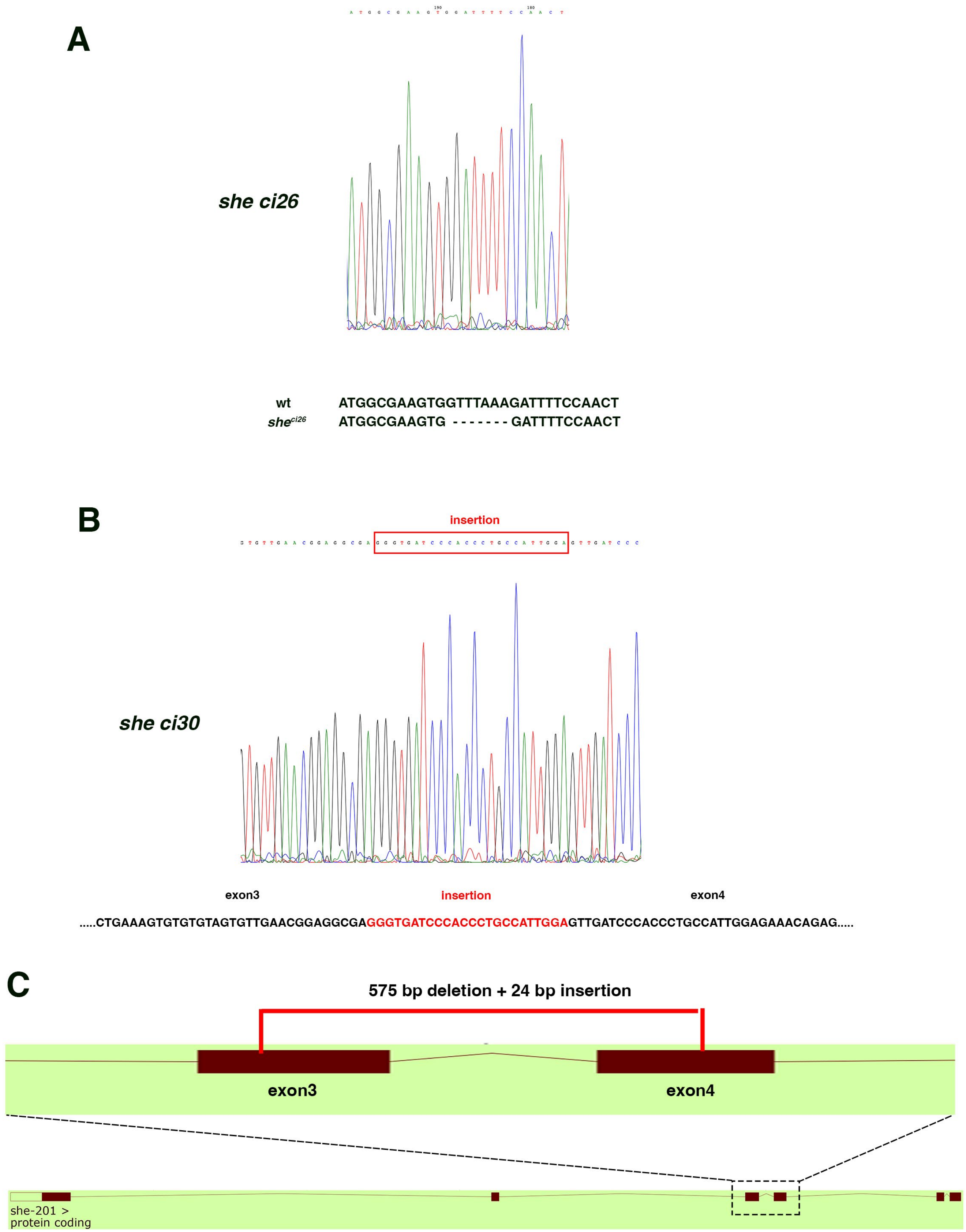


**Supplementary Figure S1. Diagrams and sequence chromatograms of *she*$^{ci26}$ and $^{ci30}$ mutations**.
(A) *she*$^{ci26}$ mutants have a 7 bp deletion. Alignment of wild-type and *she*$^{ci26}$ mutant sequences is shown starting with the first coding ATG. (B) Genomic DNA sequence chromatogram of *she*$^{ci30}$ mutants. (C) *she*$^{ci30}$ mutants carry a 575 bp deletion and 24 bp insertion present between exons 3 and 4 within *she* gene.

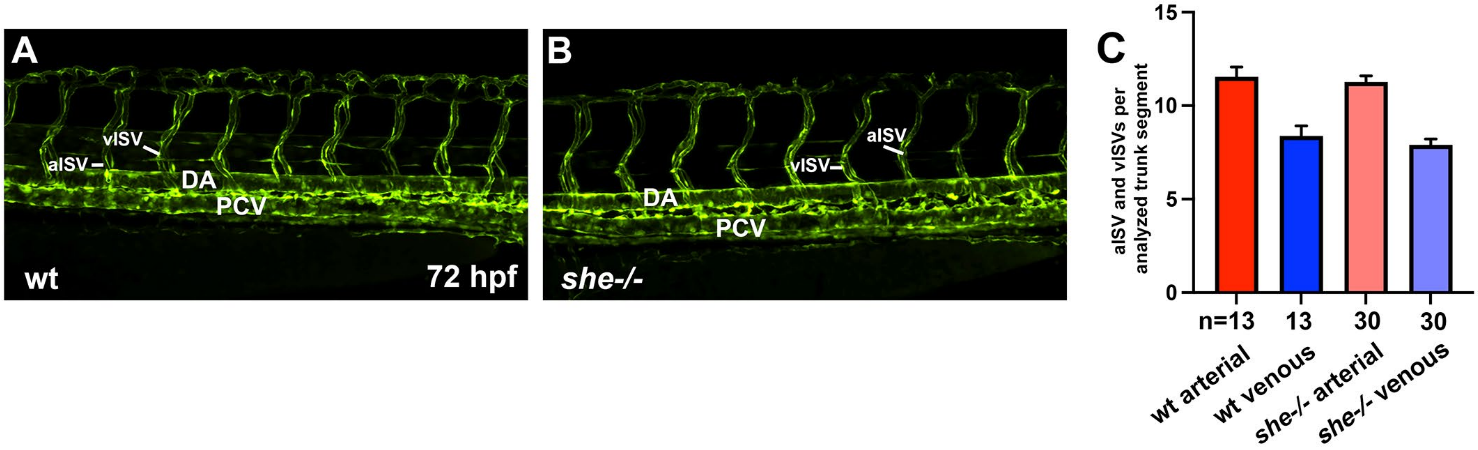


**Supplemental Figure S2. Trunk intersegmental vessel analysis in *she-/-* and wt sibling embryos.**
(A,B) Confocal images of the trunk region of live wt and *she-/-* sibling embryos in *kdrl:GFP* background at 72 hpf. Selected arterial (aISV) and venous (vISV) intersegmental vessels are shown. (C) The number of aISV and vISV was counted in a selected region in wt and *she* mutant embryos (the number of analyzed embryos in two independent experiments is provided as ‘n’). Embryos were genotyped after the imaging. No significant difference between wt and *she* mutant embryos was observed ($p>0.05$, t-Student’s test).

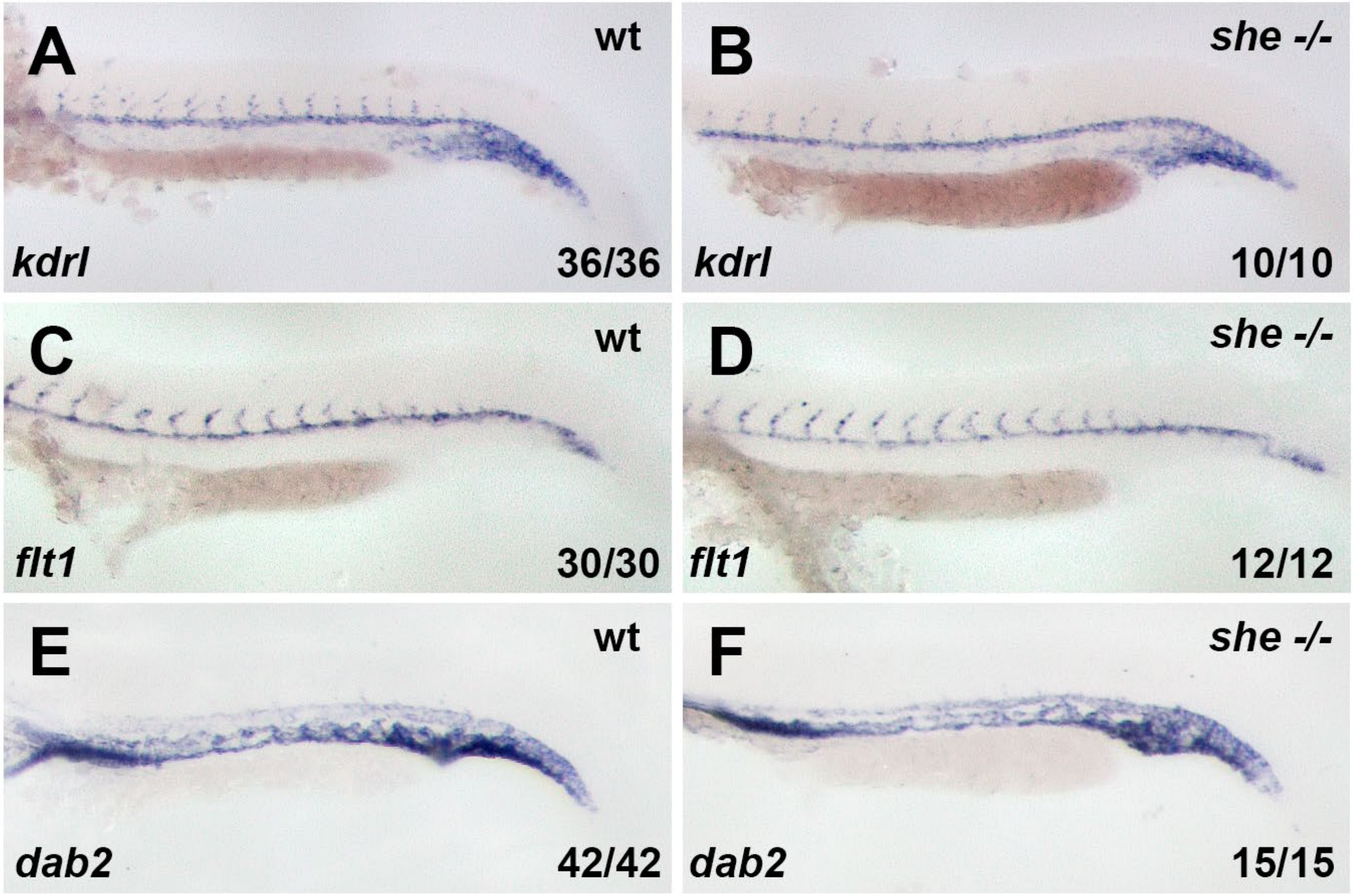


**Supplemental Figure S3. Endothelial marker expression, analyzed by in situ hybridization, is unaffected in *she* mutants at 24 hpf**.
(A, B) *kdrl* is strongly expressed in the dorsal aorta (DA) and intersegmental vessels (ISVs) and weakly expressed in the posterior cardinal vein (PCV) of both wild-type sibling and *she -/-* mutant embryos.
(C, D) *flt1* expression is restricted to the DA and ISVs of both wild-type and *she -/-* mutants.
(E, F) *dab2* is strongly expressed in the PCV and lightly expressed in the DA of both wild-type and *she -/-* mutants. Trunk region is shown in all panels. Embryos were obtained from an incross of *she+/-; kdrl:GFP* parents and subsequently genotyped.

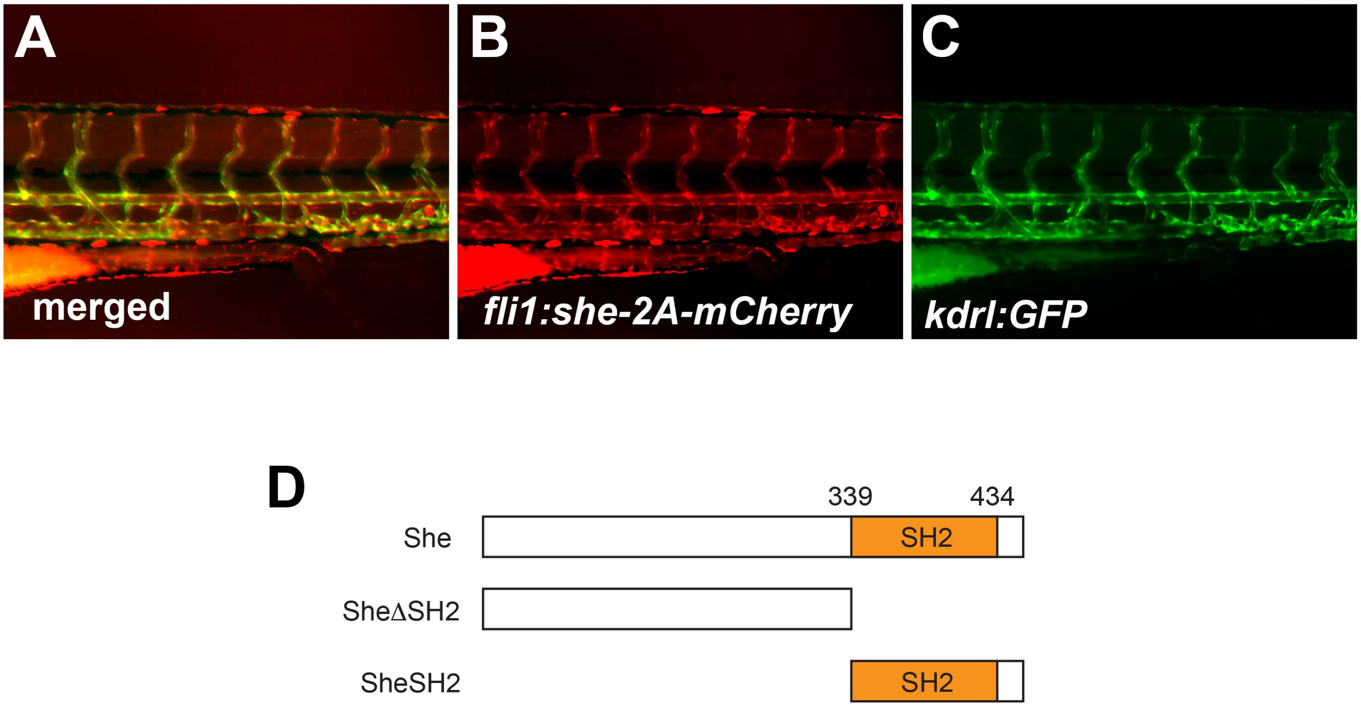


**Supplemental Figure S4. Generation of wild-type and mutant vascular endothelial-specific *she* zebrafish lines.** (A-C) Fluorescent microscopy image of *fli1:she-2A-mCherry; kdrl:GFP* embryo at 3 dpf. (D) A diagram of She deletion constructs.

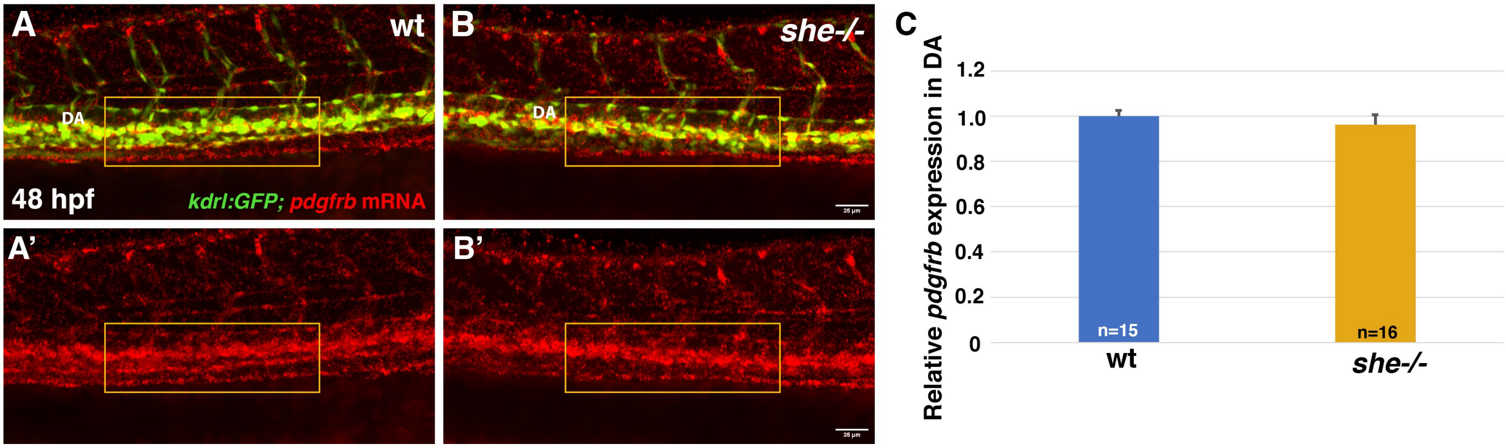


**Supplemental Figure S5.** *pdfgrb* mRNA expression analysis by hybridization chain reaction (HCR) at 48 hpf. (A,B) *pdgfrb* (red) expression in the trunk region of wild-type sibling and *she* mutant embryos in *kdrl:GFP* background at 48 hpf. Fluorescence within the dorsal aorta region (boxed) was selected for quantification. (C) Quantification of relative *pdgfrb* mRNA expression. No significant difference was observed (p>0.05, Student's t-test).

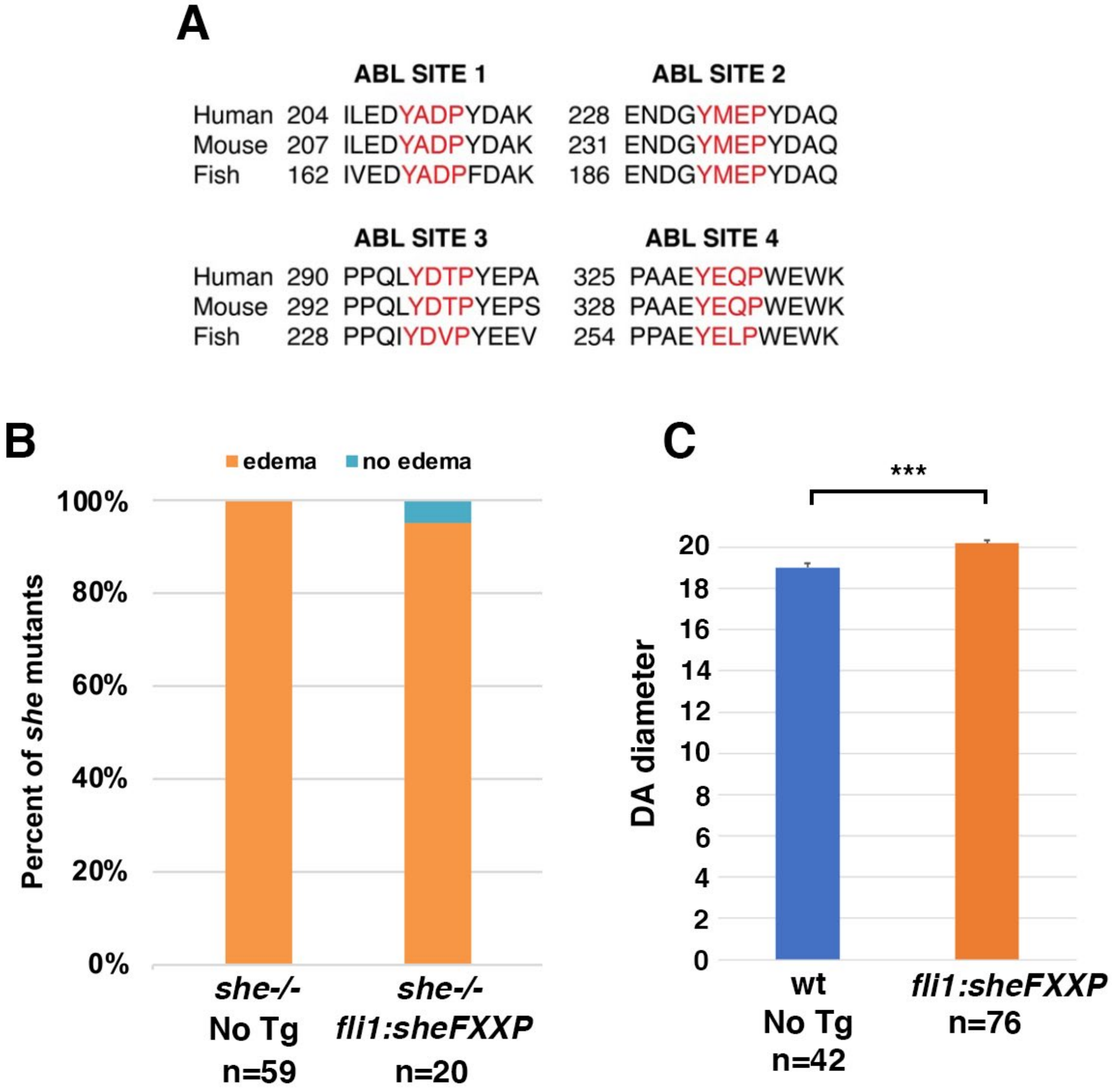


**Supplemental Figure S6. Consensus ABL phosphorylation sites YXXP are required for SHE function.**

(A) Consensus ABL phosphorylation sites are conserved between zebrafish, mouse and humans.

(B) Vascular endothelial expression of a mutant construct *fli1:sheFXXP-2A-mCherry,* where all four consensus tyrosines have been substituted into phenylalanine, fails to rescue the pericardial edema in she mutants at 4 dpf. The first bar showing *she-/-* embryos (no Tg) is copied from Fig. 3J.

(C) DA diameter measurement in *fli1:sheFXXP-2A-mCherry* embryos and sibling mCherry-negative embryos at 28 hpf. 5 measurements were performed in each embryo, which were then averaged for statistical calculations. n corresponds to the number of embryos. ± SEM is shown. ***$p<0.001$, Student's t-test.

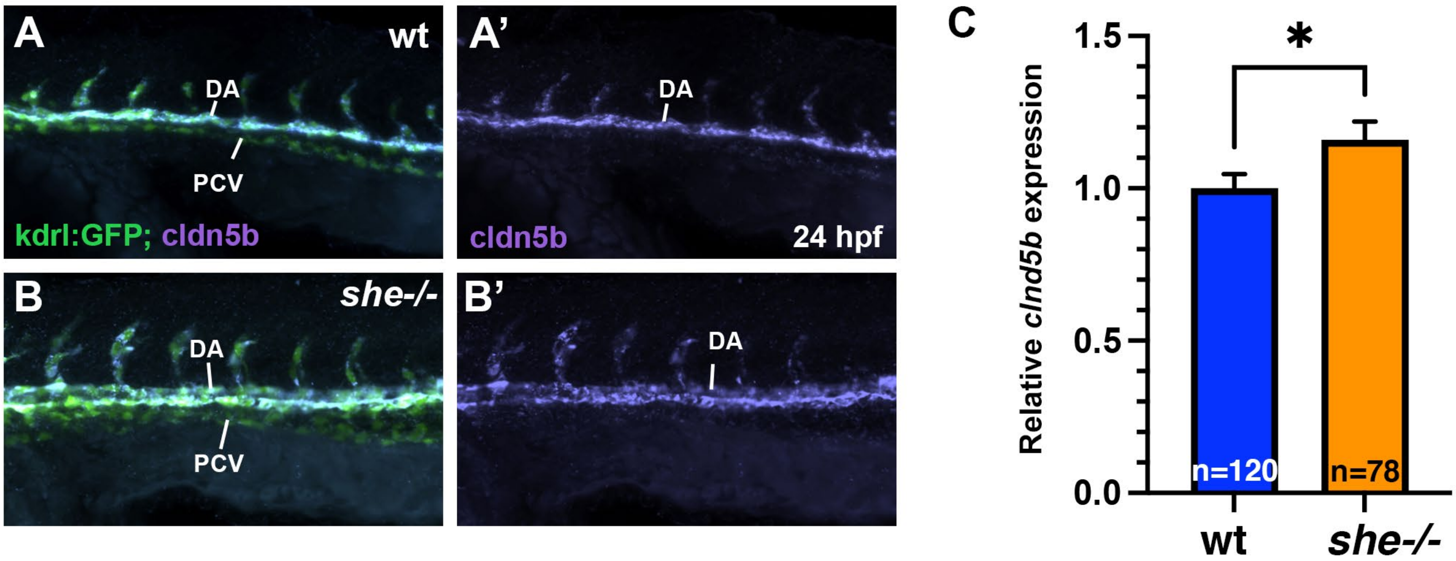


**Supplemental Figure S7. Hybridization chain reaction (HCR) analysis of *cldn5b* mRNA expression at 24 hpf.**
(A,B) *cldn5b* (purple) and *kdrl:GFP* fluorescence in the trunk region of *she* mutant and wild-type sibling embryos. DA, dorsal aorta; PCV, posterior cardinal vein. *cldn5b* fluorescence is shown in A',B'.
(C) Quantification of *cldn5b* fluorescence in the DA. *p=0.036, Student's t-test. Error bars show SEM. Data show combined results from two independent experiments.

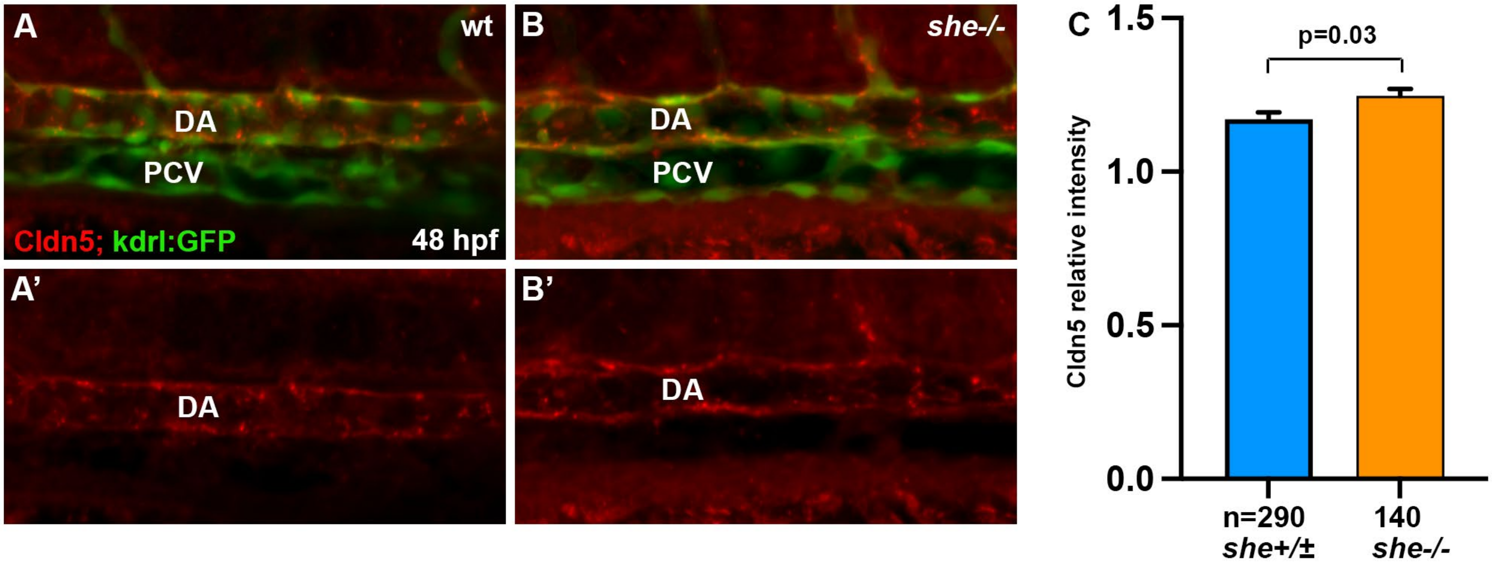


**Supplemental Figure S8. Cldn5 protein expression is increased in *she* mutant embryos.**
(A,B) Immunostaining for Cldn5 (red) combined with *kdrl:GFP* fluorescence in wild-type sibling and *she* mutant embryos at 48 hpf. Red channel is shown in A',B'. DA, dorsal aorta; PCV, posterior cardinal vein.
(C) Quantification of Cldn5 immunofluorescence within the DA. Embryos were obtained from an incross of *she+/-; kdrl:GFP* parents and subsequently genotyped. *she+/+* and *she+/-* genotypes were combined for this analysis. Error bars show SEM; Student's t-test was used.

**Supplemental Table S1. Primer sequences used to generate she mutant and transgenic lines and perform qPCR**

| | |
|---|---|
| she_F1 | ATCAATAATAGGGGCCACTTTGAG |
| she_R1 | CCGTCCTAACTCGATGGCAG |
| she_gRNA1 | GCGTAATACGACTCACTATAGGAGGCGAAGGTGAGGGGTCGTTTTAGAGCTAGAAATAG |
| she_gRNA2 | GCGTAATACGACTCACTATAGGCAGGGTGGGATCAACAATGTTTTAGAGCTAGAAATAGC |
| she_gRNA_R1 | AAAGCACCGACTCGGTGCCA |
| she_gRNA left | TATGCTCTTGTGCAGAGATTCG |
| she_gRNA right | TGACCAAAACATACCCTTTTCA |
| she_gRNA midR | CTGACAGGGCTCTAACAATCTG |
| she_attB_F1 | GGGGACAAGTTTGTACAAAAAAGCAGGCTTCACCATGGCGAAGTGGTTTAAAGATTTTCC |
| she_attB_F3 | GGGGACCACTTTGTACAAGAAAGCTGGGTTGTGTGTGCGTGGCACGGGGTG |
| she_deltaSH2_attB_R | GGGGACCACTTTGTACAAGAAAGCTGGGTTGCTCTGTTTCTCCAATGGCAGG |
| sheSH2_attB_F | GGGGACAAGTTTGTACAAAAAAGCAGGCTTCACCATGTGGTATCATGGCAGTGTGAGTCG |
| she_attB_R3 | GGGGACCACTTTGTACAAGAAAGCTGGGTTGTGTGTGCGTGGCACGGGGTG |
| sheYXXP1_F1 | CAGAACCGATCATCGTAGAGGATTTTGCCGACCCATTTGATGCTGAG |
| sheYXXP1_R1 | CTCAGCATCAAATGGGTCGGCAAAATCCTCTACGATGATCGGTTCTG |
| sheYXXP2_F1 | GAAGGAGAGAACGATGGCTTCATGGAGCCTTATGATGCCCAGCTC |
| sheYXXP2_R1 | GAGCTGGGCATCATAAGGCTCCATGAAGCCATCGTTCTCTCCTTC |
| sheYXXP3_F2 | GTCTGGACCCCCTCAGATCTTTGATGTCCCATATGAGGAGG |
| sheYXXP3_R2 | CCTCCTCATATGGGACATCAAAGATCTGAGGGGGTCCAGAC |
| sheYXXP4_F1 | CAGACCGCCGGCAGAGTTTGAGCTGCCCTGGGAGTGGAAGAAAG |
| sheYXXP4_R1 | CTTTCTTCCACTCCCAGGGCAGCTCAAACTCTGCCGGCGGTCTG |
| cldn5a-F | GACAACGTGAAAGCGCGGG |
| cldn5a-R | AGGAGCAGCAGAGTATGCTTCCC |
| EF1a-F | TCACCCTGGGAGTGAAACAGC |
| EF1a-R | ACTTGCAGGCGATGTGAGCAG |

### MOVIE LEGENDS

**Movie 1. Blood flow in the tail region of wild-type zebrafish embryos.** Wild-type siblings obtained from an incross of *she*$^{ci26}$ heterozygous adults in *kdrl:GFP; gata1:dsRed* background were imaged at 4 dpf using confocal microscopy.

**Movie 2. Blood flow in the tail region of *she* mutant embryos.** Mutant embryos obtained from an incross of *she*$^{ci26}$ heterozygous adults in *kdrl:GFP; gata1:dsRed* background were imaged at 4 dpf using confocal microscopy.
